# Accumulation of neural state transitions in dorsomedial striatum predicts patch foraging decisions

**DOI:** 10.1101/2025.09.29.679309

**Authors:** Elissa Sutlief, Shichen Zhang, Kate Forsberg, Marshall G Hussain Shuler

## Abstract

Activities with temporally distributed returns require deciding not only what to do, but when to stop. Although optimal foraging theory predicts that patch departure should depend on the total time invested in a patch, behavior across species often reflects sensitivity to recent rewards. The dorsomedial striatum (DMS) mediates both goal-directed decision-making and interval timing, two functions that converge during patch foraging, where animals must decide when to exit patches to maximize reward rates. We recorded extracellular activity from neurons in DMS while freely moving mice performed a patch-foraging task. Mice employed a reward-reset strategy, primarily basing their exit decisions on the time since the last reward, with systematic adjustments for patch residence time and environmental reward-rate context. Individual neurons in DMS underwent discrete firing rate transitions at characteristic delays following each reward. These transition times tiled the post-reward interval across the population, producing an accumulation-to-threshold signal that reset with each reward and whose rate was sensitive to the cost of time, increasing with higher environmental reward rates and as rewards occurred later in the patch. This accumulation reached a consistent threshold at the moment of patch exit, encoding the animal’s intended stopping time. Together, these results identify a time-cost-variable, reward-reset, accumulation-to-threshold computation in DMS that integrates environmental reward rate, elapsed time from pursuit engagement, and elapsed time from reward occurrence to determine when to abandon a pursuit.

## 1. Introduction

Decision-making is often framed as the selection among discrete options. However, many real- world decisions are continuous in time and instead require determining how long to persist in an ongoing activity before disengaging. This class of decisions, when to stop, is central to adaptive behavior, shaping how organisms allocate time across competing pursuits. Failures of this process are implicated in neuropsychiatric conditions such as addiction, depression, and obsessive-compulsive disorder (Addicott et al. 2017; Huys et al. 2015; Marton et al. 2019). Despite its importance, the neural mechanisms that govern the timing of disengagement from ongoing behavior remain poorly understood.

### 1.1 Ecological Models of Time Allocation

These types of continuous, temporal decisions are common within ecological foraging paradigms. Specifically, patch foraging presents animals with an environment where resources are clustered in patches. The utility of searching a patch declines over time as its resources are harvested, requiring the animal to decide when to abandon it in search of a new one. Many ethological studies have characterized various factors that affect patch occupancy time, such as patch richness (Keller & Sullivan, 2023; Marshall et al., 2013; Utsumi et al., 2009), inter-patch distance (Searle et al., 2006; Utsumi et al., 2009), predation risk (Brown, 1988; Eccard et al., 2020; Ward et al., 2000), and competition (Brown, 1988; Laguë et al., 2012). To understand how long an animal should continue searching a given patch, we use the normative principle of reward-rate maximization, which posits that the best strategy is the one that results in the highest overall rate of reward. For typical patch-foraging tasks, where the patch’s reward probability decays monotonically over time, the optimal policy is to leave a patch when the rate of reward inside the patch falls below the overall rate of reward achieved in the environment, as delineated by the Marginal Value Theorem (MVT) (Charnov, 1976). Patch occupancy across organisms is nonetheless consistently reported to exceed this optimum, in a pattern of overstaying (Baum, 1983; Bettinger & Grote, 2016; Güldener et al., 2024; Hall-McMaster et al., 2021; Harhen & Bornstein, 2023; Kilpatrick et al., 2021; Lloyd et al., 2023; Marshall et al., 2013; Nonacs, 2001; Wajnberg et al., 2000).

### 1.2 Neural Substrates of Decision-Making and Timing

The basal ganglia play a key role in action selection and option evaluation during decision- making (Cisek, 2007; Mink, 1996; Redgrave et al., 1999). The primary input nucleus of the basal ganglia, the striatum, receives cortical projections from a wide variety of regions, integrating information about current goals and shifting environmental variables to guide behavior (Balleine et al., 2007; Ferenczi et al., 2016; Frank et al., 2004). Subregions of the striatum engage with different aspects of optimizing behavior, with the ventral striatum processing reward-related information (Berridge & Robinson, 2003; Salamone & Correa, 2012), the dorsolateral striatum mediating habit formation (Packard & Knowlton, 2002; Smith & Graybiel, 2013; Yin et al., 2004), and DMS engaged in goal-directed decision-making (Balleine et al., 2007; Yin et al., 2005). Lesions of DMS severely impair behavioral flexibility in animals, causing them to lose their ability to adapt to shifting action-outcome contingencies (Castañé et al., 2010; Pisa & Cyr, 1990; Ragozzino, 2007; Torres et al., 2016; Yin et al., 2005), while electrophysiological recordings show that DMS neurons dynamically encode decision variables, including action values, predicted outcomes, and choice signals (Burton et al., 2015; Ito & Doya, 2015; Kim et al., 2013; Lau & Glimcher, 2008; Parker et al., 2016; Thorn et al., 2010).

However, much of the decision-making research in DMS uses tasks in which subjects choose between discrete options, such as two-alternative forced choice and multi-armed bandit tasks (Balleine et al., 2007; Bogacz et al., 2006; Samejima et al., 2005; Stott & Redish, 2014; Tang et al., 2022). Patch foraging presents a different type of decision where the choice measured is a continuous variable. Rather than deciding *what* action to take, subjects must determine *when* to take action. The striatum has also been strongly implicated as a region central to timing, with lesions causing impairments in interval timing and temporal discrimination (Matell & Meck, 2004; Meck, 2006; Merchant et al., 2013). DMS neurons have been shown to encode temporal information through activity patterns such as ramping (Emmons et al., 2017; Ponzi & Wickens, 2022; Vandaele et al., 2021), and temporal receptive fields (Gouvea et al., 2015; Jin et al., 2009; Mello et al., 2015). The convergent role of the DMS in both reward-related decision-making and interval timing makes it well-suited for the temporal decisions required during patch foraging.

### 1.3 Study Overview and Objectives

Here, we investigate how DMS supports decisions about when to abandon a depleting pursuit using a patch-foraging task in freely moving mice. In this task, the optimal policy depends only on total time invested in the patch, independent of individual reward times. Contrary to this prediction, we find that mice adopt a reward-reset timing strategy, in which the decision to leave is governed primarily by the time elapsed since the most recent reward, with systematic modulation by patch residence time and environmental reward rate. This strategy reflects a temporally structured process in which each reward event resets an internal timing process that determines how long the animal is willing to continue waiting.

At the neural level, we identify a corresponding population code in DMS. Individual neurons undergo discrete, reward-triggered transitions between firing-rate states at characteristic delays, collectively tiling the post-reward interval. Across the population, the number of transitioned units increases over time following each reward, forming an accumulation signal that resets with each new reward. The rate of accumulation scales with environmental reward rate and invested patch residence time, and predicts the animal’s exit timing, reaching a threshold at the moment of patch departure.

These findings reveal a mechanism by which DMS encodes a dynamically rescaled signal encoding the time to leave, integrating invested patch residence time and environmental reward rate into a unified signal that governs when to stop. More broadly, they suggest that temporal decisions to disengage may be implemented through a time-cost-variable reward-triggered accumulation process, providing a bridge between interval timing and decision-making in naturalistic settings.

## 2. Results

### 2.1 Behavioral Characterization of Patch-Foraging Strategy

We began by characterizing time investment behavior in a patch-foraging task. In some previous versions of patch foraging tasks, subjects made a series of discrete decisions to stay or leave after each reward event (Constantino & Daw, 2015; Hayden et al., 2011). Here, the patch instead delivered many randomly timed rewards, at a decreasing frequency the longer the subject stayed, so subjects self-timed their departure under a continuously decaying probability of reward, freely deciding how to invest time to maximize the overall reward rate reaped from an environment.

Mice were placed in custom behavioral chambers with access to two nose-poke ports, the ’time- investment port’ and the ’context port’ (Figure 1a). The time-investment port served as the ’patch’. Mice harvested it by holding their heads in the port and licking continuously (poke and hold).

**Figure 1:**
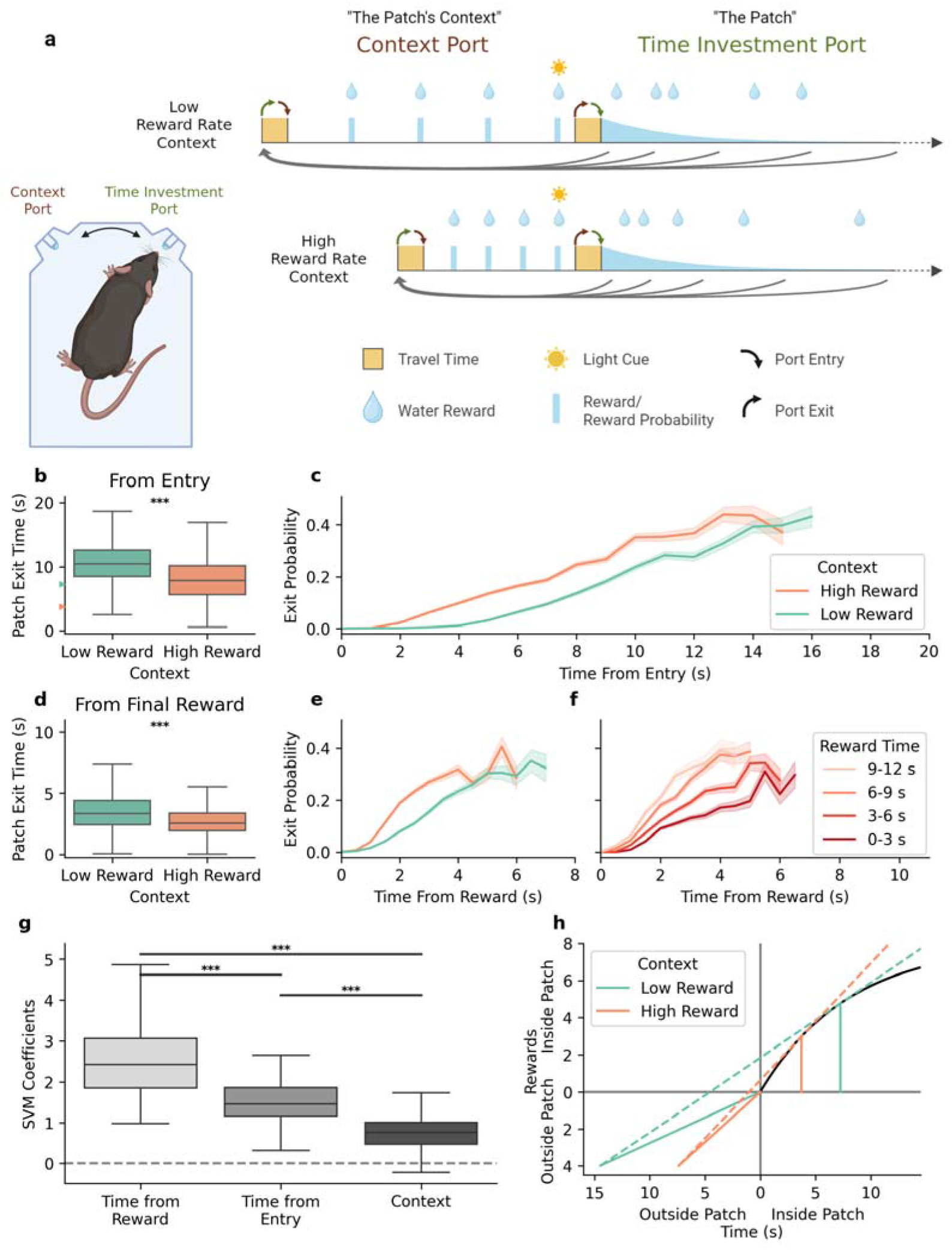
Task structure and behavioral strategy. **a)** Schematic of the behavioral box (left) and structure of the task (right). Mice alternate between collecting rewards from the “context port” and the “time-investment port”. In the context port, four rewards are delivered at regular intervals before the port becomes inactive, indicated by a light cue turning on. In the time-investment port rewards are delivered stochastically with exponentially decreasing probability. The context port is reset when the mouse enters the time-investment port, while the time-investment port is reset after the mouse receives all four rewards in the context port. Created in BioRender. Sutlief, E. (2026) https://BioRender.com/xf8ciz0. **b)** Observed exit times from the time-investment port across mice (n = 10 mice, 134 sessions, 6,970 trials), measured from the time of entry into the port (two-sample t-test, p = 1.9 × 10 ²³²). Arrows indicate the reward-rate-optimal exit times for each context (see panel h). **c)** The probability (hazard rate) of exiting the time-investment port based on the time since port entry (x-axis) and the context port reward rate (hue). **d)** Same as b, except measured from the time of the final reward received in the time-investment port (two-sample t-test, p = 8.6 × 10 ¹¹¹). **e)** The probability (hazard rate) of exiting the time-investment port based on the time since the most recent reward (x-axis) and the context port reward rate (hue). **f)** The probability (hazard rate) of exiting the time- investment port based on the time since the most recent reward (x-axis) and the time of the most recent reward relative to time of port entry (hue). **g)** Support vector machine (SVM) coefficients for predicting time-investment port occupancy. All three predictors are significantly different from zero (one-sample t-tests, all p < 0.001) and all pairwise comparisons are significant (paired t-tests, all p < 0.001). **h)** Reward-rate-optimal exit times. The origin is the moment of entry into the time-investment port. Time in the patch runs to the right and time outside it (at the context port and traveling between ports) runs to the left. Rewards earned in the patch are plotted upward and the four context-port rewards downward. The black curve is the expected cumulative reward in the patch, G(t). For each context, the solid colored line runs from the origin back through the time outside the patch to the four context-port rewards (mean across mice: 7.4 s in the high and 14.5 s in the low reward-rate context). The dashed line from that point with the highest slope that still meets G(t) gives the maximum achievable overall reward rate, and the point where it touches G(t) (solid vertical line) is the optimal exit time (3.7 s and 7.2 s, Methods).

For as long as a mouse stayed, 1 µL water rewards were delivered at random times, with a probability per unit time that decayed exponentially with time since port entry (with a time constant of 8 s, so that staying indefinitely would yield about eight rewards, see Methods). The probability of each reward was evaluated independently of other reward events, depending solely on the time since port entry. As the likelihood of reward decreased with time, mice could choose to abandon the time-investment port at any moment in favor of the context port. The context port provided a fixed number of regularly spaced rewards over a fixed interval, after which a light cue turned on to indicate that the context port was depleted and the time-investment port had been replenished. As soon as the mouse returned to the time-investment port, the light cue turned off, signaling that the context port was replenished and available, posing the challenge of deciding how much time to invest in the time-investment port. Mice learned to alternate back and forth between the two ports throughout each behavioral session, investing a self-determined amount of time in the time-investment port.

Sessions were broken into blocks where the context port switched between being ’high’ value (four rewards in five seconds) and ’low’ value (four rewards in ten seconds). The time-investment port’s reward schedule was identical in every block, so the two reward-rate contexts differed only at the context port (Figure 1a). By changing the reward rate and time spent in the context port, we altered the cost of time by manipulating the overall reward rate of the environment (Charnov, 1976; McNamara & Houston, 1985; Sutlief et al., 2025; Namboodiri et al. 2014). Changing the cost of time incentivizes reward-rate maximizing agents to leave the time-investment port earlier during high value blocks and later during low value blocks. Critically, this experimental design shifted the reward-rate optimizing patch-exit strategy solely by manipulating factors external to the patch (Figure 1h). The central behavioral measure of interest was the amount of time invested in the time-investment port during each trial. We found that, on average, mice invested significantly more time in the time-investment port during ’low’ context port blocks compared to ’high’ context port blocks (Figure 1b), progressively increasing their likelihood of departure as time passed in the time-investment port (Figure 1c). While the direction of this context- dependent shift aligns with theoretical expectations (Charnov, 1976; McNamara & Houston, 1985; Sutlief et al., 2025) and prior observations (Constantino & Daw, 2015; Kilpatrick et al., 2021; Marshall et al., 2013), mice stayed in the patch longer than is reward-rate optimal, consistent with the overstaying commonly reported in patch-foraging tasks (Charnov, 1976; McNamara & Houston, 1985; Constantino & Daw, 2015; Hayden et al., 2011; Hutchinson et al., 2008; Lenow et al., 2017; Nonacs, 2001; Wilke et al., 2009). We computed the optimal exit time for each mouse and block from that mouse’s measured time outside the patch (see Methods).

Session mean exit times exceeded the optimum by 4.5 s in the high block (one-sample t-test across sessions, t(133) = 23.5, p = 2.9 × 10 , later than optimal in all 134 sessions) and by 3.6 s in the low block (t(133) = 25.7, p = 2.0 × 10 ³, later than optimal in 133 of 134 sessions). Across sessions, mice achieved an overall reward rate of 28.7 ± 0.2 rewards/min (mean ± SEM, n = 134 sessions), with higher rates during high-context blocks (34.2 ± 0.3) than low-context blocks (23.3 ± 0.2; paired t-test, p = 1.1 × 10 ¹). This corresponded to 88.4% and 93.2% of each mouse’s optimal reward rate in the high and low blocks, respectively.

### 2.2 Time-cost–variable, Reward-Triggered Exit Time Policy

Since the probability of reward depends only on the time since entering the port, the optimal departure time is not affected by the occurrence and timing of rewards on any given trial. Nonetheless, the departure strategy exhibited by mice was very sensitive to the timing of rewards received in the time-investment port. While mice were extensively trained on the underlying reward probability distribution, the time since the most recent reward was the strongest factor determining port exit (Figure 1g), followed by patch residence time, and finally by environmental reward-rate context. Because these variables are correlated, we also measured how much each one contributed to cross-validated classification when added to or removed from the model. Time since the most recent reward alone reached a balanced accuracy of 0.90, against 0.93 for all three variables, and removing it from the full model lowered balanced accuracy the most (by 0.082, against 0.025 for time since entry and 0.013 for context, each greater than zero with p < 10 ² , n = 134 sessions). The ablation therefore gives the same order as the coefficients (Figure 1–figure supplement 4a, b). A discrete-time hazard model of leaving, which uses only the moments before each exit and so needs no ‘Leave’ labels, also gave the same order. Removing time since the most recent reward worsened the model’s cross-validated predictions the most, then time since entry, then context (paired t-tests, all p < 0.005, Figure 1–figure supplement 4d). To better understand how reward events shaped exit policy, we examined the hazard rate of leaving the patch during the interval following rewards, split by context (Figure 1e) and the time of the reward relative to port entry (Figure 1f). We found that all three of these factors contributed to exit behavior, where each reward received would "reset" the amount of time the mouse would be willing to wait for a subsequent reward prior to departure, being determined by the patch occupancy time thus far and the environmental reward-rate context. Mice thus appeared to predominantly time from the last reward, though the time they were willing to wait following each reward decreased the later the reward was received and was shorter under the high- than under the low-reward-rate context. This strategy of timing from the last reward rather than patch entry, though suboptimal, is improved by the willingness to wait progressively shorter amounts of time the longer they stay in the patch.

### 2.3 DMS Neurons Undergo Firing Rate State Transitions Following Rewards

Given that behavior was organized around reward-triggered timing, and that this timing was sensitive to both patch dwell time and time’s cost, we asked whether DMS activity expresses a corresponding representation of the post-reward interval that similarly exhibits these sensitivities, and whether that representation predicts time investment decisions.

Many DMS units showed reward-locked dynamics that resembled ramping when averaged across rewards (Figure 2c, bottom). However, inspection of single-trial responses revealed that these apparent ramps arose from discrete transitions between low and high firing-rate states following reward. This structure became evident when responses were aligned to each neuron’s inferred transition time rather than to the reward itself (Figure 2d). Figure 2a illustrates an example neuron exhibiting a reward-triggered suppression followed by activation, a pattern that recurred across the population.

**Figure 2:**
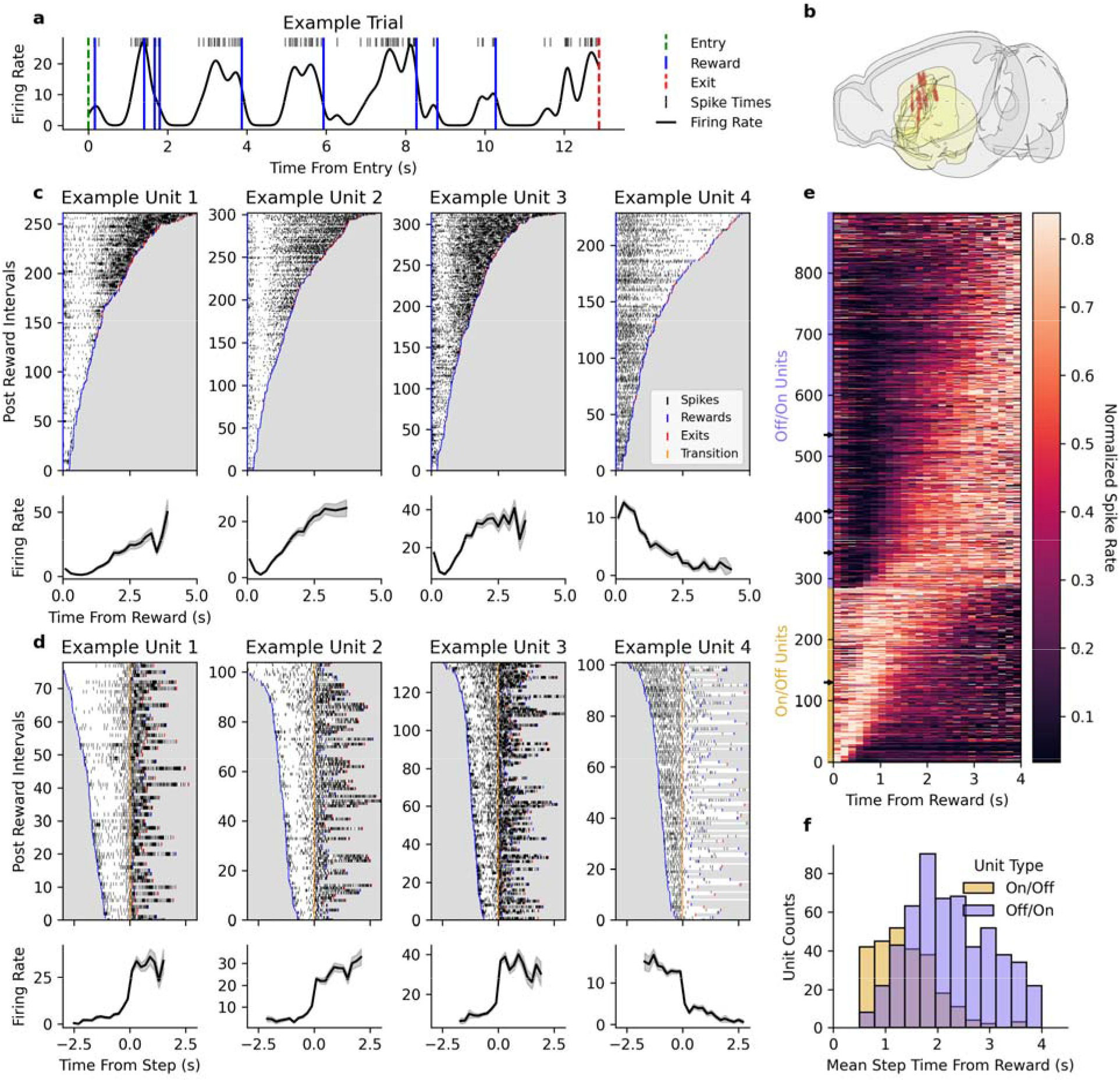
Dorsomedial striatum neurons exhibit step-like activity patterns following rewards. **a)** Example activity of a recorded unit during a single trial in the time-investment port. Top: Example unit spike raster. Bottom: Firing rate trace calculated by binning spikes (10 ms bins) and smoothed with a Gaussian kernel (σ = 120 ms). **b)** Anatomical localization of recorded units in DMS for all recorded neurons (1,849 units across 10 mice). **c)** Four example units showing reward responsive patterns. Top: Spike rasters for each reward-reward and reward-exit interval, ordered by the duration of the interval. Bottom: Corresponding mean firing-rate traces aligned to reward delivery (time = 0). **d)** Same as c, except showing intervals where each unit exhibits a state-transition, a discrete step in activity post reward, identified by the inflection point of a sigmoid function fit to the firing rate. Rasters and mean firing rate traces are aligned to the detected step times (yellow). **e)** Heatmap showing the normalized firing rates of all DMS units that make step transitions following rewards (n = 898 step-like units). Units are ordered by the direction of their activity step and their mean step time. Example units are marked with arrows. **f)** Distribution of the mean activation times of the units displayed in e.

We developed a classification procedure to identify step-like responses and estimate transition times on individual reward intervals (see Methods). Of 1,849 total units, 898 met criteria for reward-triggered state transitions (Figure 2e); validation with simulated flat-rate Poisson neurons confirmed a false positive rate of 1.35% (Figure 2–figure supplement 3). Transition times were also reproducible in data the session fit had not seen. When each unit’s session-level fit was made from a random half of its intervals, its transition time predicted the median transition time in the other half for step-like units (Spearman r = 0.73, 882 step-like units with a valid transition in each half), and much less well for matched Poisson units (r = 0.16, Figure 2–figure supplement 4). We then passed simulated neurons with continuous rate changes through the identical pipeline, at a modulation strength matched to the real step-like units (Figure 2–figure supplement 3c–h). Ramps that scaled with each interval were rarely classified as step-like (5%), and random walks in firing rate were classified as step-like in 19% of cases. Ramps with a fixed slope, which cross any given rate at the same time after every reward, usually were (92%, against 72% for simulated steps). The transitions of real step-like units were as sharp as those of simulated steps (median fitted slope 407/s and 420/s) and about four times sharper than those of the fixed-slope ramps that passed (105/s, Figure 2–figure supplement 3d).

Across these step-like units, mean transition times were distributed across the post-reward interval, effectively tiling the time axis (Figure 2f). Within units, transition times across rewards were approximately normally distributed, with variability increasing for later mean transition times (Figure 2–figure supplement 1). Apart from this reward-triggered state-switching activity pattern, several other identifiable task-related activity patterns were observed across the population, with units exhibiting, in varying combinations, responses to a variety of different task aspects, primarily to movement between ports and to licking (Figure 2–figure supplement 2). Additionally, 321 units were characterized as non-responsive and did not show any identifiable task-related activity patterns.

### 2.4 Neural State Switches Correlate with Behavioral Exit Policy

We next examined whether variability in neural transition times tracked the animal’s departure policy. Transition times covaried with several task parameters, most prominently the timing of reward relative to patch entry and context, with larger parameter correlations among units with later mean transition times (Figure 3–figure supplement 1).

**Figure 3:**
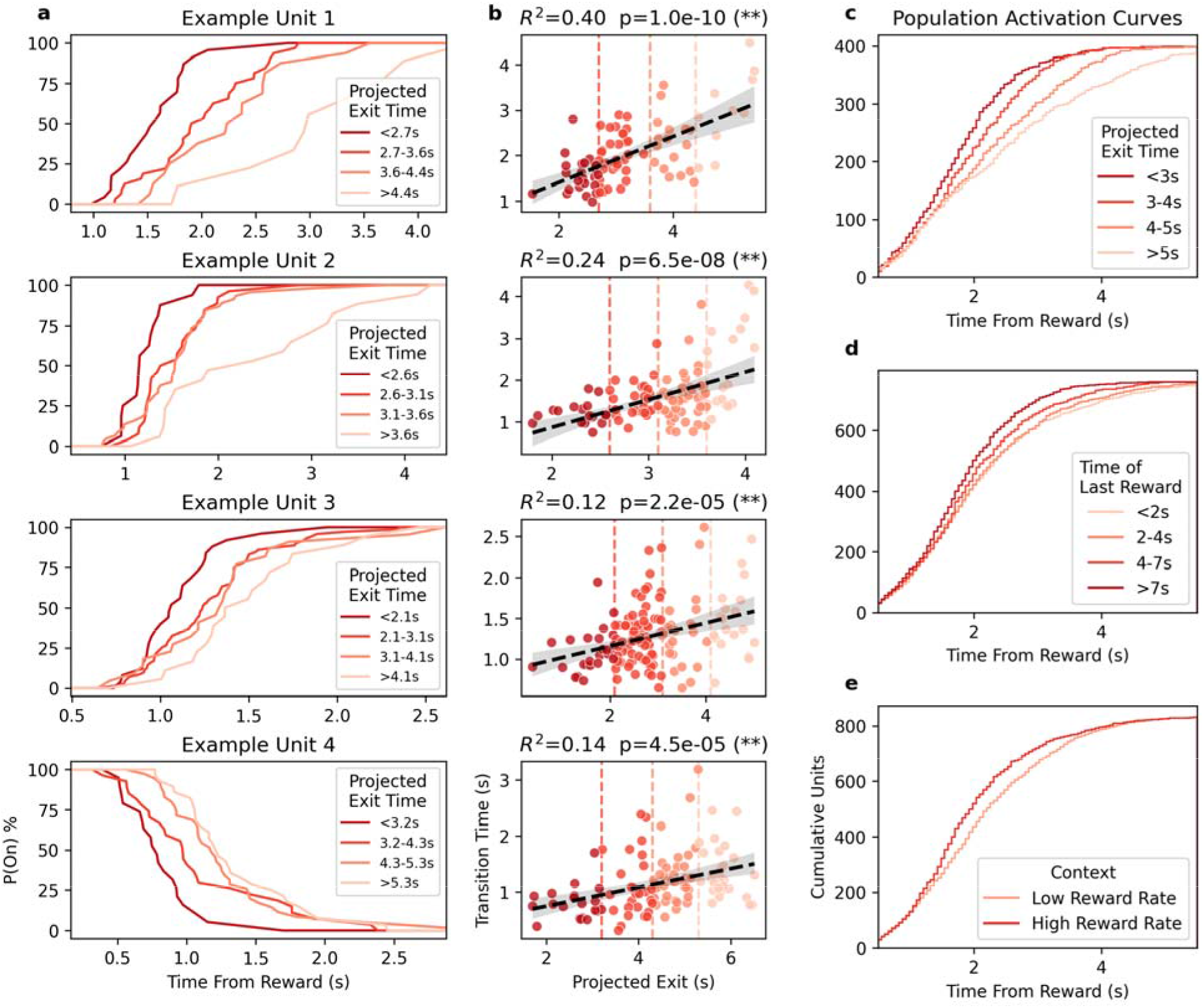
Neural state-transition times correlate with projected exit times. **a)** Plots for each example DMS unit (same as Figure 2c,d) showing the probability of the unit being in the “on” state in the time following reward, separated into four equal-width ranges of the projected behavioral exit time (see Figure 1–figure supplement 2). **b)** Correlation between unit state-transition times and projected exit times for each example unit. Hue and vertical dashed lines indicate the groupings used in a. **c)** Population mean state-transition times (n = 400 of the 898 step-like units) shown as a cumulative function for each of the four projected exit time ranges, including only units with measurements in every time range. All groups differ significantly (Kolmogorov-Smirnov, all p ≤ 0.008). **d)** Same as c (n = 762), except separated by the time of the last reward. Only comparisons involving the “>7s” group differ significantly (Kolmogorov-Smirnov; “<2s” vs “>7s”: p = 1.4 × 10 ; “2-4s” vs “>7s”: p = 3.7 × 10 ; “4-7s” vs “>7s”: p = 0.021), while remaining comparisons are not significant (all p > 0.06). **e)** Same as c (n = 840), except separated by the reward rate in the context port. Groups differ significantly (Kolmogorov-Smirnov, p = 5.3 × 10 ).

To relate neural dynamics to the mouse’s intended leave time following each reward, we derived a "projected exit time" for each reward event: an estimate of how long the mouse would have waited after that reward if a subsequent reward had not occurred (see Methods section 4.9).

Although exit timing is variable when pooling across sessions (Figure 1b), within-session policy is comparatively stable (Figure 1–figure supplement 1a). We examined several related possible predictors from the reward dynamics in the time-investment port and found that the time of the last reward relative to port entry was the most correlated with leave time from the final reward for individual sessions (Figure 1–figure supplement 1g). We therefore fitted a session-specific regression model predicting leave time from the last reward as a function of context and the time of the last reward relative to port entry (Figure 1–figure supplement 2), and used the fitted coefficients to project exit times on reward–reward intervals.

Neural transition times tracked these projected exit times. For the four example neurons in Figure 3, the probability of being in the high-firing state shifted later when the projected exit time was later and earlier when the projected exit time was earlier (Figure 3a). For each example neuron, transition time increased approximately linearly with projected exit time (Figure 3b). At the population level (Figure 3c–e), cumulative transition dynamics diverged most strongly when grouped by projected exit time: accumulation was steeper when projected exit time was earlier and shallower when projected exit time was later (all pairwise Kolmogorov-Smirnov tests significant, p ≤ 0.008; Figure 3c). This grouping produced greater separation than grouping by time since entry (Figure 3d) or by context alone (Figure 3e), consistent with projected exit time being the most behaviorally aligned regressor (Figure 3–figure supplement 1).

Step-like neurons were also present at the context port, where rewards arrive at fixed times and do not deplete, and their transitions were as sharp as those at the time-investment port (Figure 3– figure supplement 2a). Between context-port rewards, whose spacing doubles from the high to the low reward-rate block, most of these neurons transitioned at a similar latency after reward in both blocks rather than rescaling with the interval (Figure 3–figure supplement 2b), and neurons that were step-like at both ports showed little relationship between their transition times at the two ports (Figure 3–figure supplement 2c, d). Too few transitions occurred between the final context-port reward and departure, however, to test for the accumulation described below.

### 2.5 Accumulation of Neural State Transitions Predicts Exit Time

Finally, we asked whether reward-reset accumulation of state transitions provides a quantitative, trial-by-trial predictor of patch exit, and found that the rate of accumulation predicted exit timing. We analyzed the 53 sessions with at least seven simultaneously recorded step-like units (see Methods). The results below were qualitatively consistent for inclusion thresholds from 4 to 12 units (Figure 4–figure supplement 2). Figure 4a shows representative trials from an example session with 19 step-like units. Following each reward, units transitioned progressively into their post-step states, producing an accumulation of transitioned units that reset at each reward.

**Figure 4:**
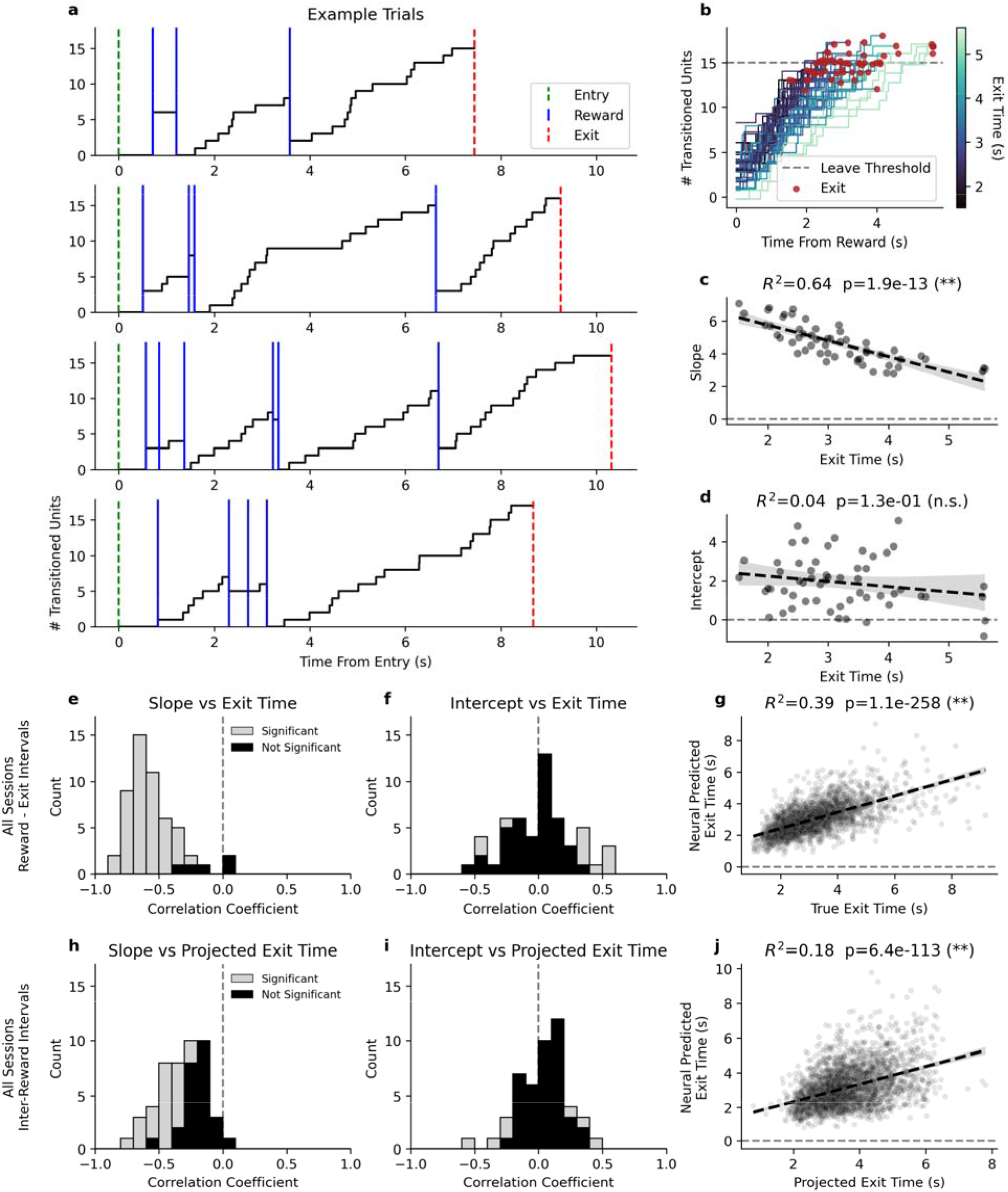
Accumulation of neural state transitions in DMS predicts exit timing. **a)** Four example trials from a single session (ES025, 56 trials, 19 step-like units), showing the cumulative step transitions of simultaneously recorded units. Vertical lines indicate entry (green), exit (red), and reward events (blue). **b)** The accumulation lines for all reward-to-exit intervals from the same session as the examples in a. Lines are colored based on the duration of time between the reward and exit events. Exits are marked by maroon dots capping each line. The gray dashed horizontal line indicates the session-specific leave threshold (median final cumulative count at exit). **c)** The slope of each accumulation line in b fit with a linear model, plotted against the exit time (R² = 0.64, p = 1.9 × 10 ¹³, OLS regression). **d)** The intercept of each accumulation line in b fit with a linear model, plotted against the exit time (R² = 0.04, p = 0.13, n.s.). **e)** Histogram of per-session Pearson correlation coefficients between accumulation slopes and exit times for all sessions with at least 7 step-like units (n = 53 sessions from 6 mice; mean r = −0.553, one-sample t- test vs. 0: t = −20.6, p = 1.0 × 10⁻²⁶; 48 of 53 sessions individually significant at p < 0.05). **f)** Histogram of per-session Pearson correlation coefficients between accumulation y-intercepts and exit times (n = 53 sessions; mean r = 0.015, t = 0.41, p = 0.68; 11 of 53 individually significant). **g)** The relationship between exit time predicted by the accumulate-to-threshold model and true exit times for all reward-to-exit intervals pooled across sessions (n = 2,410 intervals; R² = 0.388, p = 1.1 × 10 ² , OLS regression). **h)** Histogram of per-session correlation coefficients between accumulation slopes and behaviorally projected exit times during inter-reward intervals (n = 48 sessions; mean r = −0.324, t = −12.1, p = 5.2 × 10⁻¹⁶; 23 of 48 individually significant). **i)** Histogram of per-session correlation coefficients between accumulation y-intercepts and behaviorally projected exit times during inter-reward intervals (n = 48 sessions; mean r = 0.039, t = 1.37, p = 0.18). **j)** The relationship between exit time predicted by the accumulate-to- threshold model and behaviorally projected exit times for all inter-reward intervals (n = 2,622 intervals; R² = 0.177, p = 6.4 × 10 ¹¹³, OLS regression).

Across reward-to-exit intervals, accumulation dynamics scaled with exit timing: accumulation ramped faster when the mouse exited sooner and slower when the mouse exited later (Figure 4b). Notably, leading up to each exit, the accumulated count approached a similar level (15 transitioned units in this example session), consistent with a threshold mechanism. For each reward-to-exit interval, we fit a line to the accumulation trajectory and found that the slope was significantly negatively correlated with exit time (Figure 4c), whereas the intercept was not (Figure 4d).

This pattern generalized across sessions. Accumulation slopes were significantly negatively correlated with exit time (mean r = −0.553, one-sample t-test vs. 0: p = 1.0 × 10 ² , 48 of 53 sessions individually significant; Figure 4e), while intercepts showed no significant correlation (mean r = 0.015, p = 0.68; Figure 4f), consistent with a variable accumulation rate from a stable starting point. Using a threshold-crossing rule applied to accumulation trajectories, we generated neurally predicted exit times that explained 39% of variance in observed exit times (R² = 0.388, p = 1.1 × 10 ² ; Figure 4g). This prediction depended on pairing each interval’s accumulation with its own behavior. When accumulation traces were shuffled among intervals within each session, the mean per-session correlation fell from −0.553 to 0.000 (95% of shuffles between −0.044 and 0.046), the number of individually significant sessions from 48 of 53 to 2.6 on average, and the pooled R² from 0.388 to 0.028, with no shuffle reaching the matched value (10,000 shuffles, p < 10 , Figure 4–figure supplement 1). Applying the same analysis framework to inter-reward intervals (excluding reward-to-exit intervals) and comparing accumulation dynamics to behaviorally projected exit times yielded similar results (Figure 4h–j). The shuffle control gave the same result for these intervals (mean r −0.324 against 0.000 shuffled, 23 of 48 sessions significant against 2.4, pooled R² 0.177 against 0.034, p < 10 , Figure 4–figure supplement 1). We also verified that the accumulation slope was not correlated with the travel time from the investment port to the context port following exit (median 0.34 s, mean per-session r = −0.035, p = 0.22, 6 of 53 sessions individually significant, Figure 4–figure supplement 3). Together, these results identify a reward-reset accumulation-to-threshold computation in DMS that integrates elapsed time, reward history, and environmental context to predict when the animal will abandon a depleting pursuit.

## 3. Discussion

Decisions about when to disengage from an ongoing activity are fundamental to adaptive behavior, yet the neural mechanisms that govern such temporal decisions remain poorly understood. Here, we show that during patch foraging, mice adopt a time-cost-variable, reward- reset timing policy, and that this policy is reflected in a corresponding neural computation in dorsomedial striatum (DMS). Specifically, we identify a population-level signal in which discrete, reward-triggered neural state transitions accumulate over time and reach a threshold at the moment of patch exit. The rate of this accumulation varies systematically with the cost of time and predicts exit timing on a trial-by-trial basis. Together, these findings show how DMS implements an accumulation-to-threshold computation that integrates elapsed time from reward and elapsed time from pursuit initiation with environmental reward rate to determine when to depart.

### 3.1 Neural Mechanisms of Temporal Decision-Making

At the individual neuron level, we observed step-like responses in DMS neurons, differing from the ramping or sparse temporal response dynamics typically associated with timing processes. Previous studies on interval timing in the striatum have predominantly reported firing rates that increase steadily over the course of the interval (Gouvea et al., 2015) or show sparse encoding of time fields (Mello et al., 2015). At the population level, our findings indicate that the striatum exhibits activity patterns that accord with the features of the behavioral strategy employed, where the intended interval to depart from a patch appears as timed from the last reward, though modified by the context in which the patch is embedded, and by the amount of time already spent in the patch. Specifically, the number of transitioned units resets and accumulates linearly after each reward, with the slope of this accumulation predicting the actual exit time from the patch.

As with behavior, the time already spent in the patch as well as the patch context impact timing, affecting the rate of accumulation of transitioned units. When rewards occur earlier in the patch the slope of accumulation is shallower, predicting greater time investment, whereas if rewards occur later in the patch, the slope of accumulation is steeper, predicting early departure times.

When the time spent outside the patch in the context port is greater, the reward rate of the context port is lower, so that the cost of time is lower, resulting in a greater willingness to invest time in the patch. This is reflected in the correspondingly lower rates of accumulation of transitioned units in comparison to when the patch’s context is ‘high’. Collectively, these observations indicate a neural mechanism in which the DMS conveys the intended waiting interval after a reward, integrating elapsed time since last reward, elapsed time since patch entry, and the context of the patch.

### 3.2 Relationship to Accumulation-to-Bound Frameworks

The accumulation-to-threshold dynamics we observe in DMS share formal features with drift- diffusion and related accumulation-to-bound models, which have been widely applied to perceptual and value-based decisions (Shadlen & Newsome, 1996; Gold & Shadlen, 2007; Brody & Hanks, 2016). In those frameworks, noisy evidence accumulates until a bound is reached, triggering a commitment to action. Here, the accumulated quantity is not sensory evidence but the cumulative count of discrete, reward-triggered neural state transitions representing the passage of time, with the bound corresponding to the threshold at which the animal departs the patch. This parallel suggests that accumulation-to-threshold computations may serve as a general motif across decision domains, extending from perceptual discrimination to self-timed decisions about behavioral persistence. Theoretical work has explored this connection deeply (Church & Deluty, 1977; Gibbon, 1977; Gibbon et al., 1984; Killeen & Fetterman, 1988; Matell & Meck, 2000; Namboodiri et al., 2014; Simen et al., 2011; Treisman, 1963), with varying models framing interval timing as an accumulation-to-threshold process that differ mainly in the form and generation of time’s representation and how it is read-out. Additionally, models such as TOPDDM (Simen et al., 2013) and TIMERR-DDM (Namboodiri, Mihalas, & Shuler, 2016; Namboodiri & Hussain Shuler, 2016), inspired by the Behavioral Theory of Timing (BeT) (Killeen & Fetterman, 1988), adapt their timing process to the temporal structure of reward within an environment. The linear accumulation to a singular threshold we observe here accords with TOPDDM, for whereas TOPDDM encodes interval timing by adjusting the rate of linear accumulation to a threshold, TIMERR-DDM encodes interval timing by adjusting the threshold of a non-linear accumulation process. Finally, the rate-scaling we observe, in which accumulation is faster under higher environmental reward rates and longer patch residence times, is also consistent with the proposal that the average reward rate impacts the cost of time, modulating the vigor of behavioral and neural processes (Niv et al., 2007).

Several prior findings in the dorsal striatum converge with our observations. Gouvea et al. (2015) showed that striatal population dynamics predict duration judgments, and Mello et al. (2015) identified a scalable population code for time in striatum. Emmons et al. (2017) reported ramping activity in DMS during interval timing and argued that frontostriatal temporal signals are consistent with drift-diffusion models. Regarding decision-making, Yartsev et al. (2018) demonstrated a causal role for anterior dorsal striatum in evidence accumulation, and Bolkan et al. (2022) showed that DMS pathway contributions to behavior depend on cognitive demand, with strong effects when tasks require evidence accumulation. Together, these studies establish that the dorsal striatum supports both temporal processing and accumulation-dependent decision- making. Our findings extend this work by identifying, in a naturalistic foraging context, a reward-reset accumulation signal in DMS for self-timed patch departure, implemented not as continuous rate changes but as the cumulative recruitment of neurons undergoing discrete post- reward state transitions.

### 3.3 Reward-Reset Strategy and Behavioral Optimality

In contrast to the behavioral predictions of classical optimal foraging theory, we observed the use of a reward-reset strategy, where the time spent in the port before a reward predicted how long the mouse would wait following that reward. Under reward-rate maximization, the optimal patch departure time occurs when the instantaneous reward rate falls below the environmental average, as the Marginal Value Theorem states (Charnov, 1976; Kolling et al., 2012), where the timing of individual rewards in a fully stochastic environment should have no effect. However, mice consistently used the time since the most recent reward as their main decision variable, similar to the strategies previously observed in parasitic wasps (Haccou et al., 1991; van Alphen et al., 2003; Wajnberg et al., 2000). Other variables, such as the reward-rate context and patch occupancy time, had more modest but significant effects on the time to exit following each reset, demonstrating sensitivity to the time-evolving value of the patch, and to the environmental reward-rate context.

The suboptimality we observed (mice staying in patches longer than the reward-rate-optimal time) is consistent with previous reports of overstaying in patch-foraging tasks (Constantino & Daw, 2015). This pattern may reflect some inherent cognitive constraints in temporal decision- making (Cuthill et al., 1990; Kacelnik and Krebs, 1985; Krebs et al., 1974) or could represent an adaptive response to environmental uncertainty (Kacelnik and Todd, 1992; Kacelnik and Bateson, 1996; Olsson and Brown, 2006), where a longer occupancy time allows for more information collection before a certainty threshold is crossed. The reward-reset strategy may be particularly advantageous in environments where patch quality varies unpredictably, since recent rewards would provide more reliable information about the patch’s current reward rate than a previously accumulated estimate of environmental statistics. While this is notably not the case for the task described in this study, the observation that mice use a reward-reset strategy nonetheless may suggest a generalized approach to similar foraging environments.

Two questions follow from the reward-reset policy. First, what would DMS do in an animal that followed the optimal policy? In our task, the optimal exit time depends only on time since patch entry and on context. If the DMS signal follows the policy in use, we would expect accumulation in such an animal to begin at patch entry rather than reset at each reward, to ignore the timing of individual rewards, and to reach threshold at a fixed, context-specific time. In this view, the reset structure of the neural signal reflects the behavioral policy rather than a fixed property of DMS. Our data cannot test this prediction, because the mice did not follow the optimal policy. Second, would a reset policy, and a reset neural signal, also appear when gain is continuous? Our rewards are discrete equal packets delivered at random times from a continuously decaying probability.

The optimum is computed from the expected cumulative reward, a smooth curve, while the animal experiences a stochastic staircase. Our data cannot answer this, since every reward in our task was a discrete packet.

### 3.4 Limitations and Future Directions

Our findings described here are correlational. Follow-up causal experiments could further support these conclusions by suppressing DMS activity during patch occupancy. If the accumulation signal we describe sets the animal’s intended departure time, then such suppression should not simply shift exit times in one direction but should degrade their structure. Exit times should become more variable once context and the time of the last reward are taken into account, and should lose their dependence on both. This would extend findings in interval-timing tasks, where striatal inactivation reduces rats’ sensitivity to duration (Gouvea et al., 2015) and disrupting either D1 or D2 neurons in DMS slows mice’s timed responses (Bruce et al., 2025). A second possible outcome is that the animal disengages from the task, which would be harder to interpret. Motor effects of DMS suppression would need to be measured in either case, for example from the time taken to travel between ports. Timing and movement can be separated. In Bruce et al. (2025), the disruption lengthened timed responses without changing how long mice took to move between nosepokes, and slowing striatal dynamics by cooling stretches timing judgments without a matching effect on movement (Monteiro et al., 2023). The result that would falsify our account is exit timing that stays as predictable, and as sensitive to context and reward history, under DMS suppression as without it. Our data also cannot distinguish a signal generated in DMS from one inherited from cortical inputs such as orbitofrontal or medial prefrontal cortex, with DMS acting as a relay. Timing-related ramping in DMS has previously been shown to depend on medial frontal cortex in rats (Emmons et al., 2017).

How downstream circuits read a count of transitioned neurons is not known. One possibility is that structures downstream of the striatum in the basal ganglia carry the same transitions. The accumulation could then sharpen along this chain, from the linear rise we observe in DMS to a more threshold-like switch closer to motor output, which would trigger departure. Whether a transition count is carried and sharpened along this pathway remains to be tested. Finally, we saw no relationship between the prevalence of step-like neurons and probe placement within DMS, although our placements were not designed to sample the structure evenly.

### 3.5 Conclusion

In summary, we identify a neural mechanism in which discrete, reward-triggered state transitions in DMS accumulate to a threshold that determines when to abandon a pursuit. This time-cost- variable, reward-reset accumulation provides a parsimonious account of how temporal information, reward history, and environmental context are integrated to guide decisions about when to stop. We suggest that DMS plays a central role in generating the temporal signals that underlie time-investment strategies during foraging, with downstream circuits likely converting this timing representation into the motor action to depart. Our findings are correlational, and causal manipulations will be needed to determine whether the accumulation dynamics we observe are necessary for patch departure decisions or instead reflect timing signals inherited from cortical or other inputs to DMS. Additionally, the generality of the reward-reset accumulation mechanism across different foraging environments and task structures remains to be tested. Nonetheless, by linking behavioral strategy to a specific population-level neural computation, these findings bridge interval timing and decision-making, and offer a framework for understanding how the brain controls persistence and disengagement in dynamic environments.

## 4. Methods

### Key Resources Table

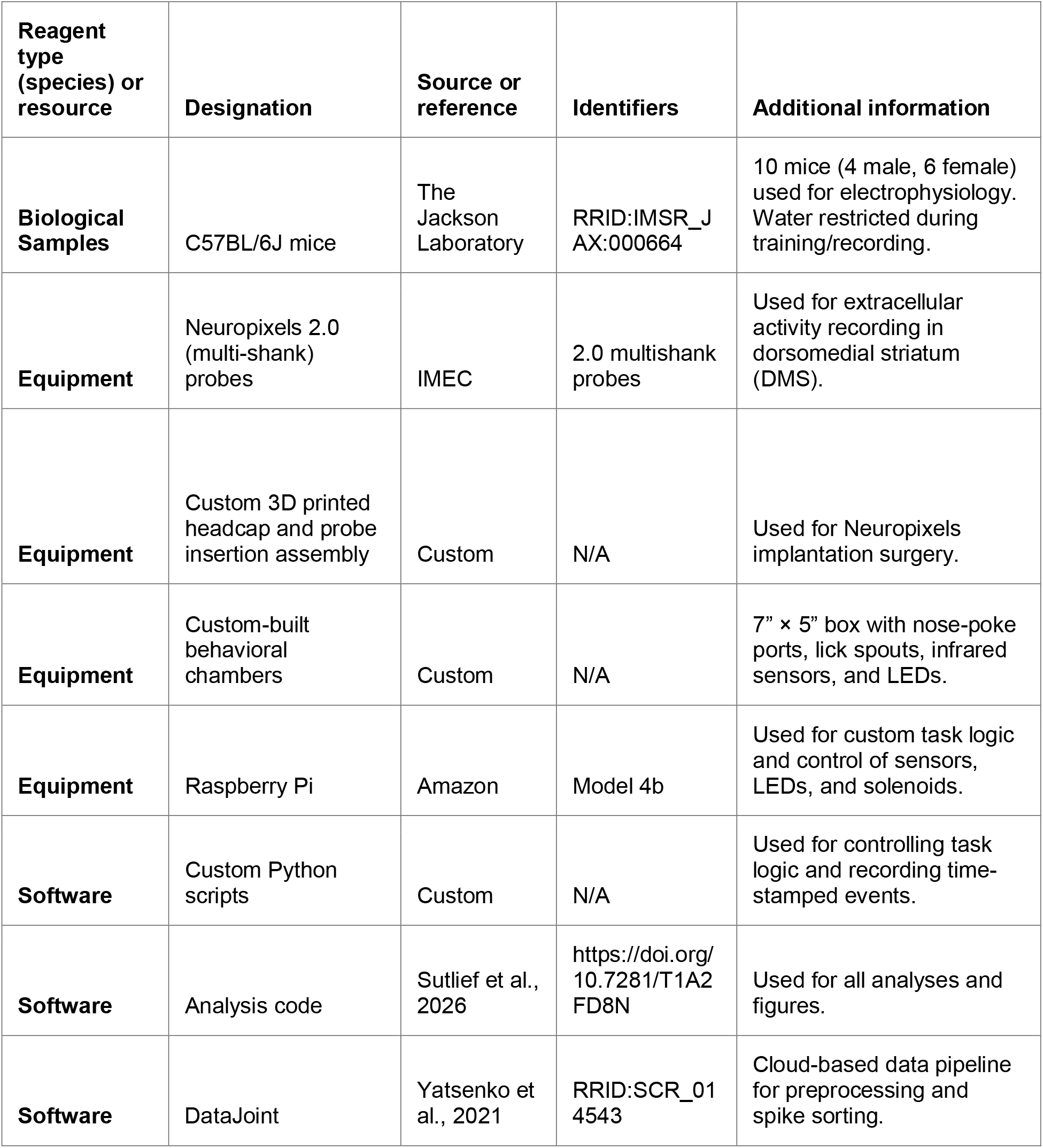

| **Reagent type (species) or resource** | **Designation** | **Source or reference** | **Identifiers** | **Additional information** |
|---|---|---|---|---|
| **Biological Samples** | C57BL/6J mice | The Jackson Laboratory | RRID:IMSR_JAX:000664 | 10 mice (4 male, 6 female) used for electrophysiology. Water restricted during training/recording. |
| **Equipment** | Neuropixels 2.0 (multi-shank) probes | IMEC | 2.0 multishank probes | Used for extracellular activity recording in dorsomedial striatum (DMS). |
| **Equipment** | Custom 3D printed headcap and probe insertion assembly | Custom | N/A | Used for Neuropixels implantation surgery. |
| **Equipment** | Custom-built behavioral chambers | Custom | N/A | 7” × 5” box with nose-poke ports, lick spouts, infrared sensors, and LEDs. |
| **Equipment** | Raspberry Pi | Amazon | Model 4b | Used for custom task logic and control of sensors, LEDs, and solenoids. |
| **Software** | Custom Python scripts | Custom | N/A | Used for controlling task logic and recording time-stamped events. |
| **Software** | Analysis code | Sutlief et al., 2026 | https://doi.org/10.7281/T1A2FD8N | Used for all analyses and figures. |
| **Software** | DataJoint | Yatsenko et al., 2021 | RRID:SCR_014543 | Cloud-based data pipeline for preprocessing and spike sorting. |

### 4.1 Compliance Statement

All animal procedures were conducted in accordance with the National Institutes of Health guidelines and were approved by the Johns Hopkins School of Medicine Institutional Animal Care and Use Committee (IACUC).

### 4.2 Subjects

Ten C57BL/6J mice (4 male, 6 female) were used in this study. Mice were purchased from The Jackson Laboratory and housed in standard vivarium cages. All animals had ad libitum access to food and water, except during periods of behavioral training and electrophysiological recording, when water was restricted. All animals maintained 85-90% of their original body weight throughout water restriction.

### 4.3 Task Design

Mice were trained on a patch-foraging task designed to specifically observe and measure the moment when a mouse decides to give up on a depleting patch (Figure 1a). The timing of that decision optimally depends on the relative expected value of remaining in the patch compared with the overall expected value of the environment. In our task, mice are required to harvest from a patch under low and high reward-rate environments while the underlying reward probability function within the patch remains consistent. Broadly, the task requires the mouse to switch back and forth between two ports as it harvests water rewards. The two ports are the time-investment port, which functions as a patch with reward delivered stochastically according to an exponentially decaying distribution, and the context port, which delivers four evenly spaced rewards over either a short or a long interval, depending on block.

The task is performed in a custom-built 7” × 5” box where the mouse has access to two ports into which it may poke its head. The ports are equipped with lick spouts that deliver 1 µL water rewards, infrared sensors that detect head entry and licks, and LED light indicators that signal whether or not the port is active (LED off = port is active/rewards are available, LED on = port inactive/rewards are not available). The mouse begins in the first of the two ports, the context port, where it receives 4 water rewards regularly spaced over either 5 or 10 seconds, depending on the current block. After the final reward is delivered, the context port light turns on and the time-investment port light turns off, indicating the mouse may then switch ports. Travel between the ports takes approximately 300 ms. Upon entering the time- investment port, the context port light turns off, signaling that the mouse may abandon the time- investment port in favor of the predictable context port rewards at any time. For as long as the mouse chooses to stay, the time-investment port delivers rewards probabilistically, with the likelihood of reward decreasing exponentially as time passes. Specifically, rewards are delivered at a rate λ(t) = exp(−t/τ) per second, where τ = 8 s and t is the time since port entry. Reward delivery is evaluated every 100 ms, with a probability of λ(t) × 0.1 s at each evaluation, so the first evaluation after port entry has a probability of 0.1. The expected number of rewards after time t in the port is approximately G(t) = τ(1 − exp(−t/τ)), which approaches eight for indefinite occupancy, while the actual number delivered is unbounded. The rate depends only on time since port entry and is the same in every block. Eventually, the moment of experimental interest arrives, being when the mouse decides to exit the time-investment port and return to the context port. As soon as it reenters the context port, the time-investment port light turns on and a single cycle (trial) is complete. Each time the mouse consumes a set of four context port rewards, the time-investment port reward probability is refreshed to its starting value, allowing the mouse to continuously harvest by alternating between the ports.

Mice learned to operate the task without the need for any shaping steps. To consider a mouse fully trained, we required that they did not leave the context port prematurely on more than 20% of the trials and that they move to the time-investment port within 3 seconds of the light cue turning on in the context port. This ensured that they experienced the intended reward rate when deciding when to leave the time- investment port. Mice typically reached criterion within one to two months of daily training (two sessions per day). No requirements are placed on their strategy in the time-investment port.

Each behavior session lasted 18 minutes and was divided into six three-minute blocks. Across blocks, the dynamics of the context port alternated between providing a high reward-rate context (4 rewards in 5 seconds) and a low reward-rate context (4 rewards in 10 seconds). The value of the starting block (‘low’ or ‘high’) was randomly determined for each session. The reward probability distribution of the time- investment port remained consistent throughout all trials across all mice. All of the logic of the task, as well as control of the sensors, LEDs, and solenoids, was custom-written in Python and deployed on a Raspberry Pi. All time-stamped events were saved to a text file for later analysis.

Exit was defined as the moment the mouse’s head left the time-investment port’s infrared beam for the last time before it entered the context port. Mice could leave the time-investment port and return without visiting the context port. Absences shorter than 2 s were treated as part of the visit. Trials in which the mouse left for 2 s or more and returned, or took 2 s or more to reach the context port, were excluded. These rules excluded 3.2% of trials. The time from exit to context-port entry was measured from the same beam events.

### 4.4 Analysis - Support Vector Machine (SVM) Model

To understand the factors contributing to the decision of when to leave the time-investment port, a Support Vector Machine (SVM) classifier was used to identify the conditions under which a mouse would either stay or leave. Three factors were supplied to the SVM: The time since entry into the time- investment port, the time since the most recent reward in the time-investment port, and the reward-rate context (either low or high). Each trial from a given session was divided into 100 ms time bins and assigned to the ‘Stay’ group. The five seconds following each exit was similarly divided and assigned to the ‘Leave’ group. Time since entry and time since the most recent reward were measured from the time- investment port throughout, including in the five seconds after exit. Labeling these seconds as ‘Leave’ rests on an assumption. Because the patch’s reward rate only declines with time, a moment after the animal chose to leave offers conditions no better than those it had just rejected, so it is treated as a moment at which the animal would also leave. Without these samples each trial would contribute a single ‘Leave’ point, leaving the decision boundary poorly constrained. The SVM then used a linear kernel to calculate the plane that best separated the two groups. The coefficients of each normalized feature were then compared across sessions to determine the relative importance of the time from reward, the time from entry, and the reward-rate context to the patch-leave decision strategies used by mice across sessions. To test whether the results depend on the length of the post-exit window, we refit the model in every session with windows of 1, 2, 5, 10, 20 and 100 s. The order of the coefficients was the same at every window (time since the most recent reward against time since entry, paired t-tests, p = 0.018 at 1 s and p < 10 at every longer window). Because the features are z-scored over all samples, including the post-exit ones, the coefficients’ magnitudes change with the window, so we also compared the decision boundary itself, expressed as the projected time from a reward to leaving, for rewards 2, 5 or 10 s after entry in each context. Its median change from the 5 s window was under 0.1 s for windows of 2 to 20 s, and at most 0.47 s at 1 s and 0.33 s at 100 s (Figure 1–figure supplement 4c).

To assess each feature’s contribution without relying on coefficient magnitudes, we fit the same classifier to every non-empty subset of the three features in each session. Performance was measured by three-fold cross-validated balanced accuracy (the mean of the correct-classification rates for ‘Stay’ and ‘Leave’), with folds grouped by interval so that no interval contributed to both training and testing. For each feature, the drop in balanced accuracy when it was removed from the full model measured its unique contribution. Drops were compared with paired t-tests across sessions (n = 134), with Wilcoxon signed- rank tests as a check.

As a complementary analysis that needs no ‘Leave’ labels, we fit a discrete-time hazard model of leaving in each session. This was a logistic regression on 0.1 s bins from port entry up to and including the bin containing the exit, with outcome 1 in the exit bin and 0 in every earlier bin, the same three z-scored predictors and the same trials as the SVM. Each predictor’s unique contribution was the rise in three-fold cross-validated log loss, with folds grouped by trial, when it was removed from the full model. Contributions were compared with paired t-tests across sessions (n = 134, all models converged).

### 4.5 Surgery for Neuropixels Implantation

Mice underwent stereotaxic surgery to implant Neuropixels 2.0 (multi-shank) probes using a custom 3D printed headcap and probe insertion assembly. All surgeries were performed in aseptic conditions and followed an IACUC approved protocol. Animals were anesthetized with isoflurane and their body temperature was maintained at 37°C. Ophthalmic ointment was applied to the eyes and Lidocaine (0.0021 mL/g) was injected subcutaneously under the scalp. Fur was removed from the scalp before it was disinfected with alternating Betadine and 70% isopropyl alcohol. A midline incision exposed the skull, with excess skin removed to expose a large surface area of skull to ensure the headcap could be securely attached. The periosteum was cleared with alternating 3% hydrogen peroxide and 70% isopropyl alcohol. The skull was then leveled and the craniotomy target (dorsomedial striatum, AP: 0.7 mm, ML: 1.2 mm) was measured from bregma. The skull surface was thoroughly dried and a thin layer of Metabond dental cement was applied to the entire exposed surface, which was given 10 minutes to fully set. The craniotomy target was then marked on the Metabond, and an oval section (approximately 1.2 mm × 1 mm) was drilled through both Metabond and skull. A separate small hole was drilled for a ground pin posteriorly on the contralateral side. A small section of dura was carefully removed from the craniotomy. The custom headcap was then placed over the craniotomy, with an oval lip fitted inside to prevent skull regrowth where the Neuropixels probe would be inserted. A gold grounding pin was placed in the posterior hole. Metabond was used to secure both headcap and pin to the skull, ensuring the craniotomy and ground pin hole were fully enclosed. Buprenorphine (0.0083 mL/g) was then administered subcutaneously on the lower back, approximately 5 minutes prior to the end of the surgery. The Neuropixels 2.0 probe was then slowly inserted into DMS, using a built-in screw and shuttle system custom designed to interface with the headcap. The probe was lowered to a target depth of 5 mm at a rate of 1 mm/min. The ground pin was connected to the grounding pad of the Neuropixels probe via a soldered silver wire and the Neuropixels housing assembly was secured to the headcap with two external screws. The weight of the entire piece reached approximately 3 grams. Mice were supplied with hydration gel and monitored as they recovered in a heated cage until they were fully awake.

### 4.6 Electrophysiology

Neural activity was recorded from DMS of fully trained mice using Neuropixels 2.0 multi-shank probes. Each probe has four 10 mm shanks, with a total of 5120 recording sites across them. The acquisition system used included a National Instruments PXIe-1071 chassis, with a PXIe-8381 module. Neural signals were recorded with a sampling rate of 30 kHz across 384 channels using SpikeGLX software.

Before recording, activity across the probe shanks was observed. The surface of the brain was located and used to estimate the band of recording sites most likely to cover the DMS, which the 384 channels were then assigned from which to record. Surface row information was saved to later validate histologically determined unit locations.

Probes were implanted using custom 3D printed fixtures for chronic, freely moving recordings, and the locations of all identified units were verified *post hoc* (Figure 2b). After recovery from surgical probe implantation, mice were retrained and DMS activity recorded for up to a month, depending on stability. Recordings were preprocessed using built-in methods from SpikeInterface (Buccino et al., 2020) and sorted using a consensus-based algorithm combining Kilosort 2.5 and Kilosort 3 (Pachitariu et al., 2016), all of which were orchestrated and managed with the DataJoint infrastructure (Yatsenko et al., 2021).

### 4.7 Histology

At the end of the experiment, mice were anesthetized with isoflurane and injected with sodium pentobarbital (200 mg/kg). Mice were then transcardially perfused with phosphate-buffered saline (PBS), then 4% paraformaldehyde (PFA). Brains were removed and placed in 4% PFA overnight, then transferred to PBS. Brains were sliced coronally at 50 µm using a vibratome (Leica VT1000 S), then mounted on glass slides. Prior to surgery, the probes were coated in Vybrant DiI (V22885) and the tracks were then imaged using a fluorescence microscope (Keyence BZ-X800). Probe track locations were reconstructed within the Allen Mouse Brain Atlas using HERBS (https://github.com/Whitlock-Group/HERBS).

### 4.8 Neural Data Preprocessing and Spike Sorting

Recording sessions were processed using a cloud-based pipeline implemented with DataJoint (Yatsenko et al., 2021) and running on AWS cloud services. Code for the preprocessing and sorting steps is publicly available at https://github.com/dj-sciops/jhu_shuler-lab_element-array-ephys.

Preprocessing was conducted using the SpikeInterface toolbox. First, a bandpass filter was applied to the raw signals to remove frequencies below 300 Hz and above 6000 Hz. Broken channels were identified and interpolated over so they would not disrupt the spike sorting algorithm. Phase shift correction was applied to account for the phase delays between channels during the system’s acquisition cycles. Lastly, common average referencing (CAR) was applied across all channels to reduce common noise artifacts.

Spike sorting was performed independently with Kilosort 2.5 and Kilosort 3. Because the two algorithms complement each other in detecting different signal types, we used the SpikeInterface consensus method to identify units that were reliably detected by both sorters, ensuring that only high-confidence units were carried forward. Quality metrics were calculated for all consensus units, then assessed during manual curation performed using Phy, where remaining noise units were removed. The final set of curated units for each session was stored in the cloud pipeline and made available for analysis. Software versions for all preprocessing, spike sorting, and analysis pipelines are specified in the accompanying code repositories (see Data Availability Statement). Statistical analyses were performed in Python 3.12 using SciPy 1.17.0, NumPy 1.26.4, scikit-learn 1.8.0, and statsmodels 0.14.6.

### 4.9 Analysis - Behaviorally Projected Exit Time Calculation

To predict how long a given mouse on a given session would be willing to wait following each reward, we fit the leave time following each final reward (the dependent variable) as a function of when that reward occurred relative to port entry (the predictor), using simple linear regression.

Separate models were fit to trials from each context block to allow for shifts in both the slope and intercept of the correlation. We then used the regression models to predict how long following every reward the mouse would have been willing to wait if no further rewards had been delivered on that trial.

### 4.10 Analysis - Neural Step-like Classification

To characterize neurons exhibiting step-like responses at varying temporal delays from reward, an algorithm was developed to identify high and low firing rate states across trials and precisely detect the timing of transitions between states. To do so, first the instantaneous firing rate of individual neurons was calculated as one over the interspike interval for 10 ms time bins. For each unit, all of the post-reward responses in the time-investment port were pooled together and a sigmoid curve was fitted to the unsmoothed firing rate data. The parameters (y-offset, height, slope, and center) of this session sigmoid were then used as initial parameters for sigmoid fits to every interval individually. Parameter bounds were set to force the individual interval (reward-to-reward and reward-to-exit) sigmoid fits to match the high and low firing-rate states identified in the session sigmoid fit. The absolute value of the sigmoid slope was required to be greater than 11 (in units of delta(spikes/second)/second), preventing the sigmoid from fitting ramping activity with shallow curves. The center parameter (inflection point of the sigmoid) was bounded to the duration of the interval plus or minus 10%. The primary criterion for a successful fit was for the sigmoid to fit the center point within the interval (at least 100 ms after the start and 100 ms before the end). On intervals where there was no clear step, the sigmoid’s best fit pushed the center to one of the far edges of the interval, essentially fitting the interval as entirely low firing or entirely high firing, due to the strict bounds on the other parameters. Because intervals could be shorter than a given unit’s typical transition latency, classification was restricted to intervals long enough for a transition to plausibly occur. Specifically, units were taken as clear examples of step-like units if the algorithm successfully identified a step on at least 50% of intervals whose duration exceeded the unit’s mean transition time plus one standard deviation of its transition time distribution, with at least 10 detected steps and a mean transition time of at least 0.5 s after reward. A linear support vector machine separating these examples from the other units set a boundary in the plane of the coefficient of variation (CV) of each unit’s transition times and its Matthews correlation coefficient (MCC), which measures how well a firing-rate threshold separates 250 ms bins before and after its transitions. Units above the boundary were classified as step- like. To validate this pipeline, matched homogeneous Poisson process neurons (flat-rate, no step structure) were generated for each recorded unit and processed through the identical classification procedure, yielding a false positive rate of 1.35% (95% CI [0.85%, 2.04%]; Figure 2–figure supplement 3).

To test whether detected transitions depend on the session-level fit, each unit’s reward intervals were split at random into two halves. The session-level fit was made from one half only, and the intervals of the other half were fit under its bounds, each fit starting from the held-out half’s own session-level transition time, so that the only information passed from one half to the other was the bounds. We compared the session fit’s transition time with the median transition time of the held-out half, with the roles of the halves swapped and both directions kept, for units with at least one valid transition in each half (Spearman correlation). The same procedure was applied to Poisson neurons matched to each step-like unit’s firing rate and intervals.

To test continuous alternatives, we simulated four alternative activity patterns for each real step-like unit, each with that unit’s intervals and firing-rate states. In the interval-scaled ramp, the rate rose from the low to the high state across each interval, whatever its length. In the fixed-slope ramp, the rate rose with the same slope in every interval, spanning the two states over the unit’s median interval, and was cut off at the interval’s end. In the random walk, the rate started at the low state after each reward and drifted at random with no trend, with a spread over the median interval equal to the difference between the states. As a positive control, the step switched abruptly between the states at a time drawn from the real unit’s transition times. The modulation was set to half the fitted difference between the states, the strength at which simulated steps matched real step-like units in Matthews correlation coefficient, transition-time variability and transition sharpness. Simulated spike trains went through the identical classification pipeline. Transition sharpness was the fitted sigmoid slope averaged over a unit’s valid intervals.

At the context port, steps were detected with the same procedure on the context-port reward intervals, and units were classified with the same boundary as at the time-investment port. The accumulation analysis used the same inclusion criteria as in Figure 4. Each port’s analysis included the units with at least one usable interval at that port, so the totals differ slightly (1,890 units at the context port and 1,849 at the time-investment port, 1,833 in both). For inter-reward intervals, each unit’s median transition time was compared between blocks for units with at least five valid transitions in each block. The ratio of transition times between the blocks was estimated with a linear mixed model of the logarithm of each transition time, with block as a fixed effect and a random intercept and random block effect for each unit (208 units, 15,041 transitions). The exponentiated block coefficient is the ratio of low-block to high-block transition times (1 if transitions keep the same latency after reward, about 2 if they rescale with the interval). Chance overlap between ports was computed within ten groups of session firing rate.

### 4.11 Analysis - Neural Accumulation

Sessions with at least seven simultaneously recorded step-like units, not counting units that stayed in their post-transition state through 70% or more of their reward intervals, were included in the accumulation analysis. This minimum was chosen to ensure sufficient population sampling for reliable linear regression of cumulative transition counts. Results were qualitatively consistent across inclusion thresholds from 4 to 12 step-like units (Figure 4–figure supplement 2). Thresholds of 4 or fewer are equivalent, because fitting an interval’s accumulation line requires at least five step positions. The cumulative number of units in their post-transition state was measured over the course of each trial. The slope and intercept of the accumulation across each reward-reward and reward-exit interval was calculated using simple linear regression. To predict the leave time from the neural activity, a single cumulative unit threshold per session was calculated as the median number of transitioned units at the time of each true exit within that session, after excluding exits more than two standard deviations from that median. This approach was selected after evaluating several threshold-setting methods; a single session-wide median threshold sufficiently captured the accumulation-to-threshold pattern observed in the data. The predicted leave time for each interval was then calculated as the intersection of its regression line with the session threshold.

To test whether the prediction depends on the pairing of accumulation with behavior on each trial, the accumulation traces (slope and intercept together) were shuffled among intervals of the same type within each session, with behavior and the session’s threshold held fixed, and the per-session correlations, the number of significant sessions and the pooled R² were recomputed (10,000 shuffles). p-values are the fraction of shuffles at least as far from the shuffle mean as the matched value. The accumulation slope of each reward-to-exit interval was also correlated, per session, with the time from exit to context-port entry on the same trial, using the same session criteria as Figure 4e. All tests were two-sided.

### 4.12 Analysis - Reward-Rate-Optimal Exit Time

The optimal exit time was defined as the time in the patch that maximizes the overall reward rate across a full cycle of the task (Sutlief et al., 2025). A cycle consists of a visit to the time-investment port lasting t seconds, which yields G(t) rewards in expectation, and the time outside it, T, which yields the context port’s four rewards. The overall reward rate is R(t) = (G(t) + 4) / (t + T), and the optimal exit time t* is the value of t that maximizes it, found numerically. At the maximum, the instantaneous reward rate in the patch equals the overall rate, λ(t*) = R(t*). This is the condition of the Marginal Value Theorem (Charnov, 1976), extended to a task in which the time between patch visits is itself rewarded. Two features of our task differ from the classical setting, and the calculation handles both directly. Rewards are discrete and stochastic, so G(t) is the expected cumulative reward. Time outside the patch also earns reward at the context port, which enters the numerator. For each mouse and block, T was the mouse’s mean time at the context port in that block plus its mean travel time from the patch to the context port and back, giving optimal exit times of 3.2 to 5.0 s in the high block and 6.4 to 8.4 s in the low block. Figure 1h shows the construction for the mean time outside across mice (7.4 s in the high block and 14.5 s in the low block), which gives optimal exit times of 3.7 s and 7.2 s.

To test whether mice stayed longer than optimal, we took each session’s mean exit time in each block over all patch visits (7,490 visits in 134 sessions), subtracted the mouse’s optimal exit time for that block, and compared the differences with zero (two-sided one-sample t-test). Efficiency was each session’s reward rate in each block, total rewards over total time, as a fraction of the mouse’s optimal reward rate R(t*) for that block.

## Data Availability Statement

The data and analysis code are available in the Johns Hopkins Research Data Repository at https://doi.org/10.7281/T1A2FD8N (Sutlief et al., 2026).

## Author Contributions (CRediT)

Elissa Sutlief (First Author): Conceptualization, Investigation, Data Curation, Formal Analysis, Methodology, Project Administration, Software, Visualization, Writing – Original Draft, Writing – Review & Editing Shichen Zhang (Author 2): Investigation, Methodology, Formal Analysis, Writing – Review & Editing Kate Forsberg (Author 3): Investigation, Methodology Marshall G Hussain Shuler (Senior Author / PI): Conceptualization, Supervision, Funding Acquisition, Project Administration, Writing – Review & Editing

## Acknowledgments

We thank Dr. Patricia Janak, Dr. Dan O’Connor, and Dr. Jeremiah Cohen for project guidance and valuable comments on the manuscript. We are also grateful to the DataJoint team for technical support. We also thank Rebekah Zhang, Dr. Simon Allard, and Dr. Tanya Martin, and the rest of the Hussain Shuler lab for their support.

---

**Figure 1–figure supplement 1:**
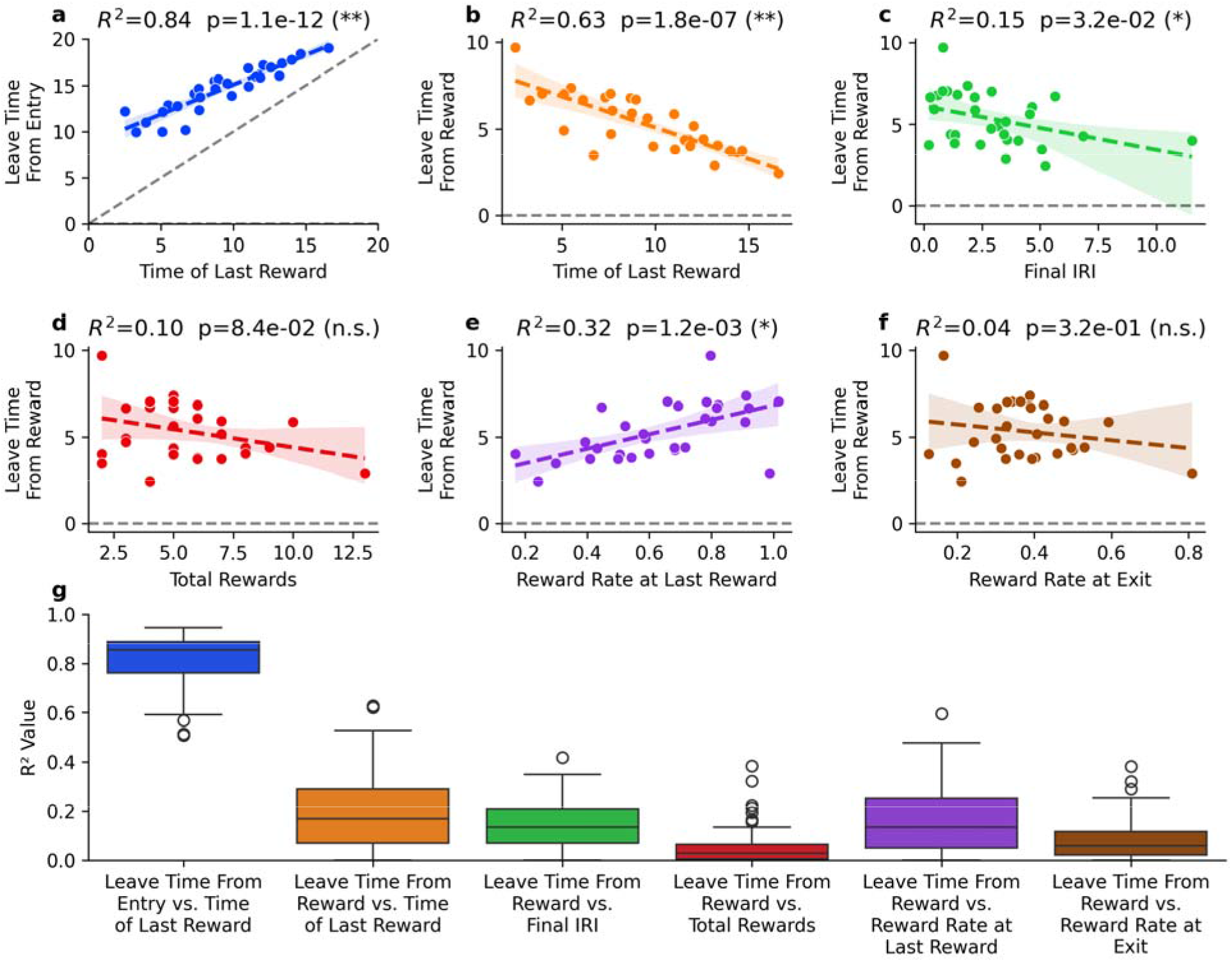
Session-specific behavioral correlates of patch exit timing. a-f) Linear regression analysis of potential predictors of patch exit behavior from an example session. **a)** Leave time from port entry plotted against the time of the last reward relative to entry. **b)** Leave time from the final reward plotted against the time of the last reward relative to entry. **c)** Leave time from the final reward plotted against the duration of the fina inter-reward interval. **d)** Leave time from the final reward plotted against the total number of rewards received in the time-investment port during that trial. **e)** Leave time from the final reward plotted against the reward rate experienced in the time between port entry and the final reward. **f)** Leave time from the final reward plotted against the reward rate experienced in the time between patch entry and patch exit. **g)** Distribution of correlation strengths (R²) across all recording sessions for each of the six behavioral predictors analyzed in panels **a-f**. Each data point represents the R² value from the linear regression for a single session. Box plots show median, quartiles, and range of correlation strengths.

**Figure 1–figure supplement 2:**
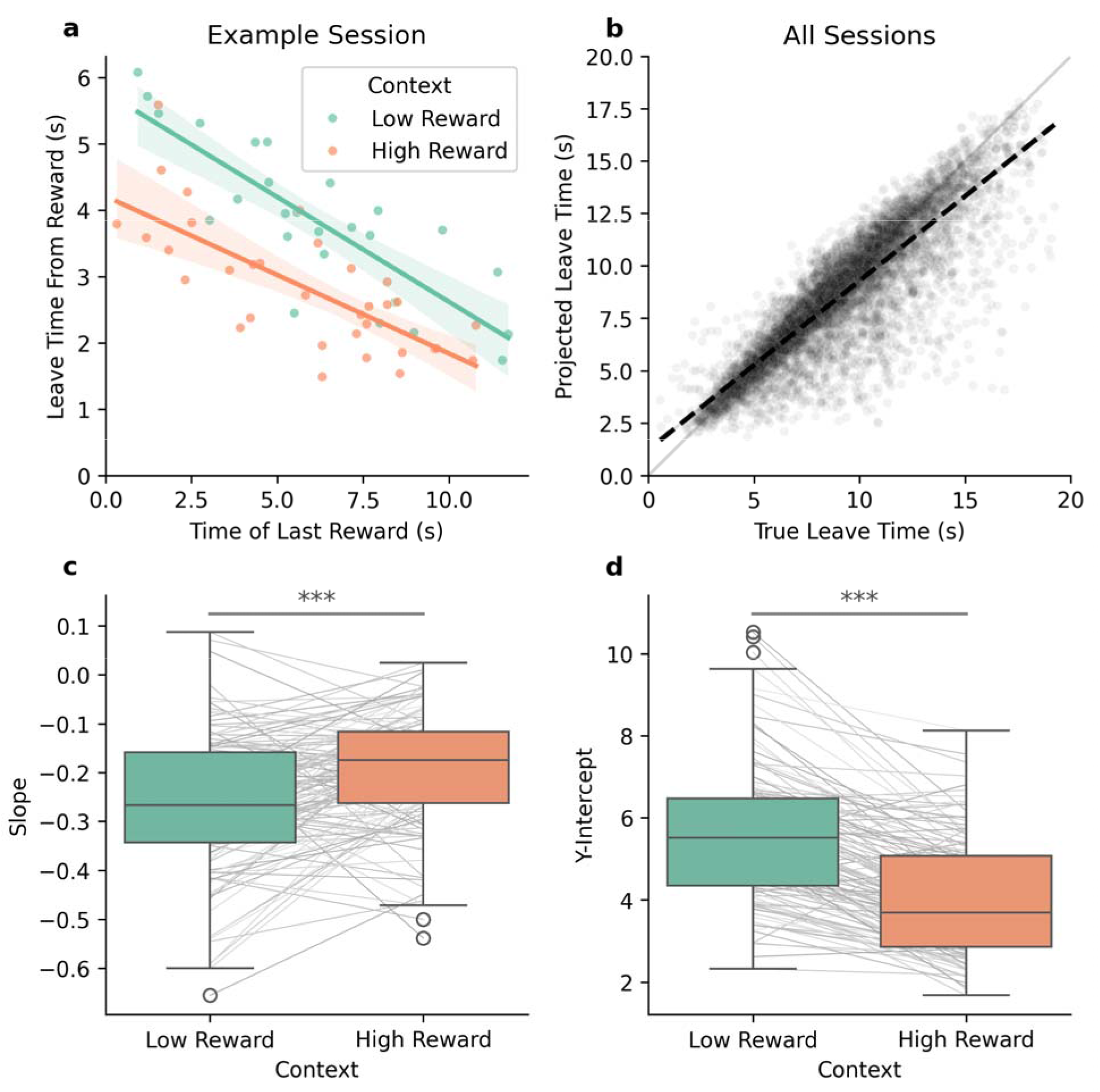
Session-specific linear regression models predict behavioral exit policy following rewards. **a)** Leave time from the final reward plotted against the time of the final reward relative to port entry for an example session. Data points are separated by context block reward rate with fitted linear regression lines. **b)** Projected leave times from port entry calculated using session-specific linear regression coefficients plotted against observed leave times across all sessions (R² = 0.75). **c)** Comparison of regression line slopes between high and low reward-rate context blocks across sessions (n = 134 sessions, p = 8.5×10^-8^, paired t-test). **d)** Comparison of regression line intercepts between high and low reward-rate context blocks across sessions (n = 134 sessions, p = 2.6×10^-27^, paired t-test).

**Figure 1–figure supplement 3.**
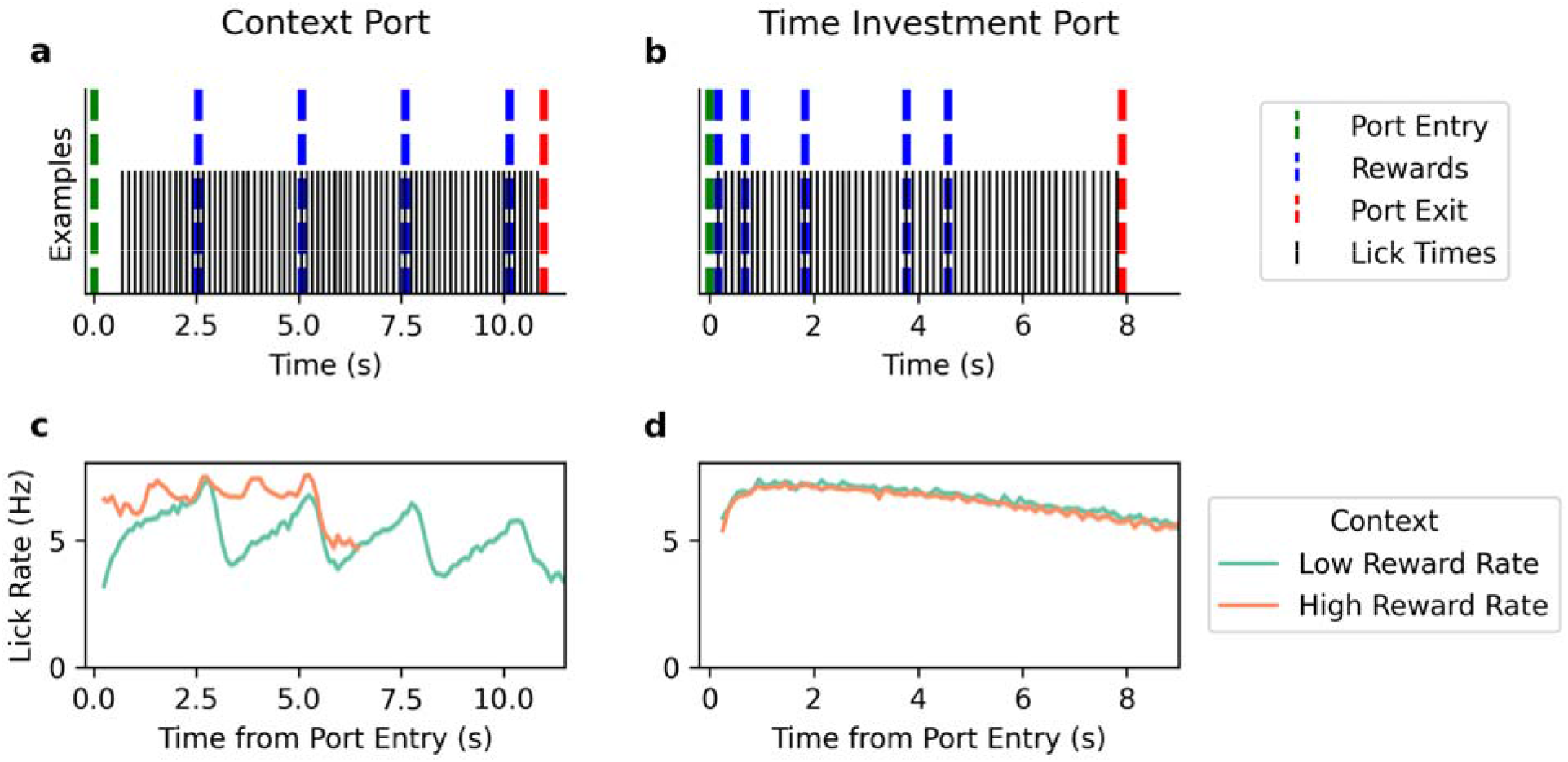
Lick patterns at the context and time-investment ports. **a)** Example single-trial lick pattern during a visit to the context port. Black tick marks indicate individual lick times. Dashed vertical lines mark port entry (green), reward deliveries (blue), and port exit (red). **b)** Same as **a** for a visit to the time-investment port. **c)** Mean lick rate as a function of time from port entry at the context port, plotted separately for low (green) and high (orange) reward-rate contexts. Shading indicates ±1 SEM across intervals. **d)** Same as **c** for the time-investment port. At both ports, licking begins shortly after entry and is sustained throughout the visit. At the context port, lick rate is higher and more sustained in the high reward-rate context.

**Figure 1–figure supplement 4.**
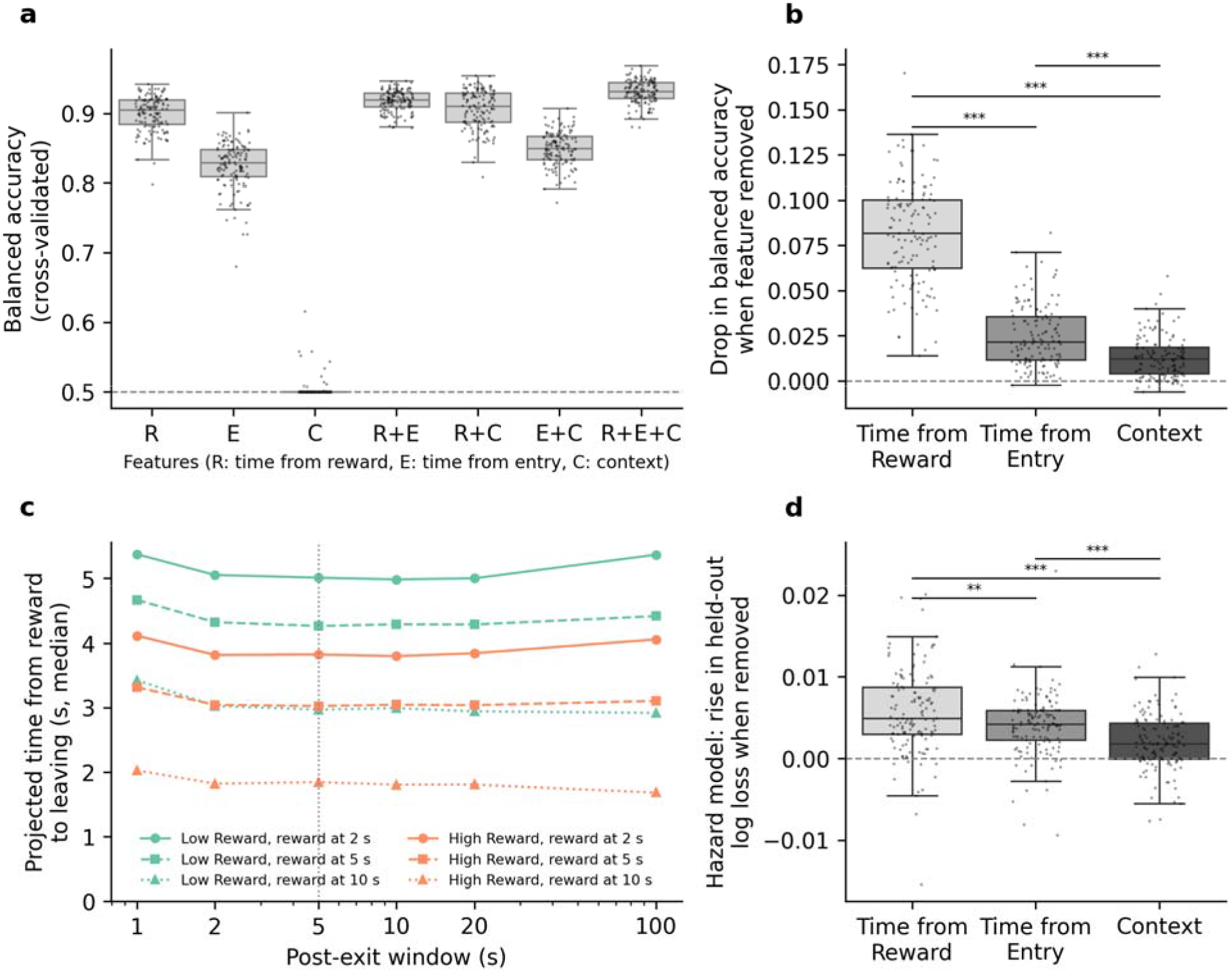
The contribution of each behavioral variable to the leave decision. **a)** Cross- validated balanced accuracy of the support vector machine of Figure 1g refit with every subset of its three features, per session. R stands for time since the most recent reward, E for time since entry and C for context. Each interval between entry, rewards and exit was kept within a single fold. Dashed line, chance. **b)** Drop in balanced accuracy when each feature is removed from the full model. **c)** The Figure 1g decision boundary expressed in seconds, as the projected time from a reward to leaving, for rewards 2, 5 or 10 s after entry in each context, across post-exit windows of 1 to 100 s (medians across sessions). Dotted line, the 5 s window used in Figure 1g. **d)** The rise in cross-validated log loss when each predictor is removed from a discrete-time hazard model of leaving (logistic regression on 0.1 s bins up to the exit, no post-exit samples), per session. In a, b and d, box plots show median, quartiles and range, and points are sessions (n = 134). Paired t-tests across sessions, *** p < 0.001, ** p < 0.01.

**Figure 2–figure supplement 1:**
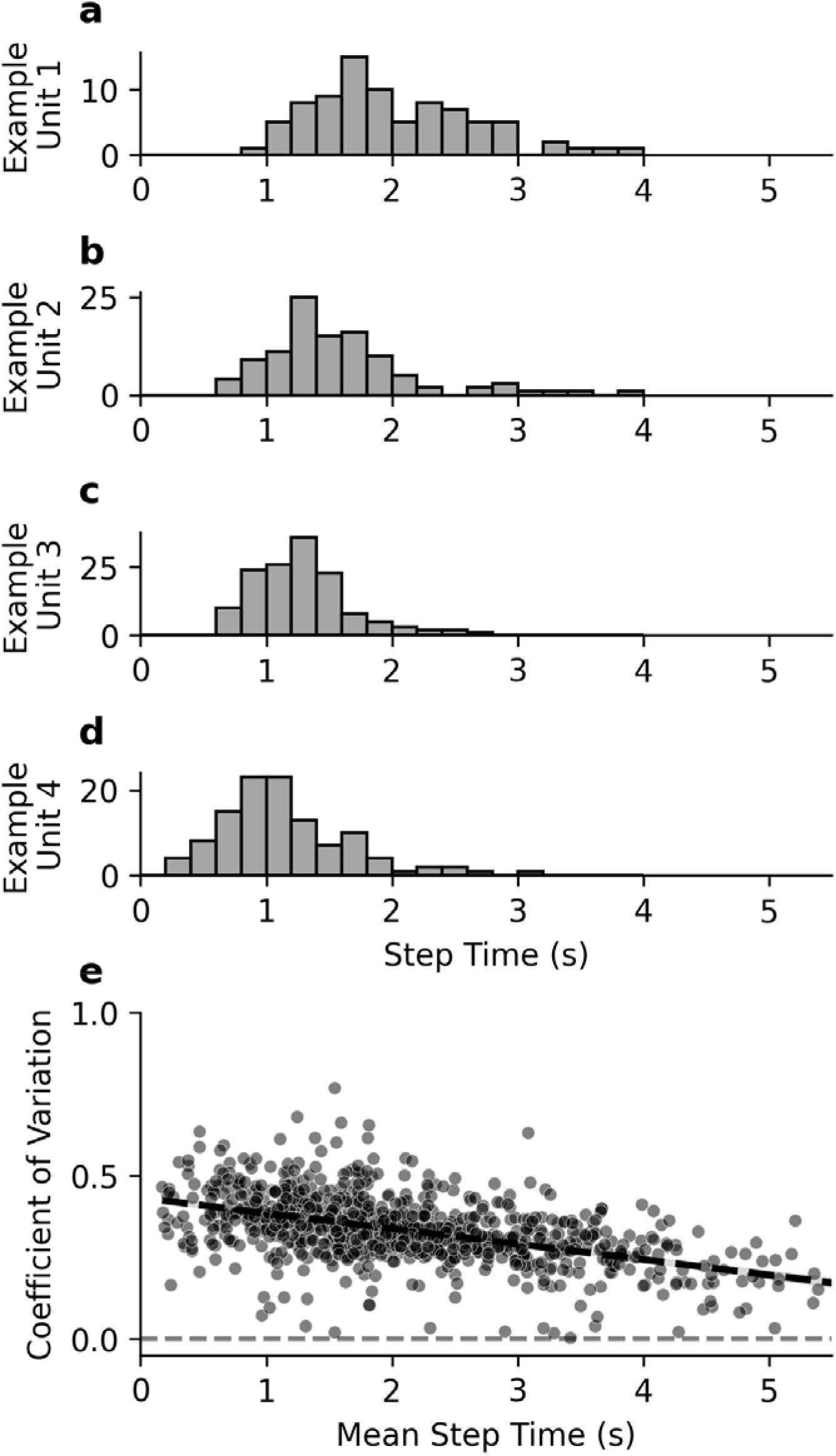
Step time distributions and variability scaling in DMS neurons. a-d) Histograms showing the distribution of step times for the four example DMS units from Figures 2 and 3. **e)** Relationship between the mean step time and coefficient of variation (CV) of step times across all step-like units (n = 898 units). Linear regression line is shown in black (R² = 0.281, p = 2.7 × 10 , OLS regression). Later-stepping units tend to have lower temporal variability relative to their mean step time.

**Figure 2–figure supplement 2:**
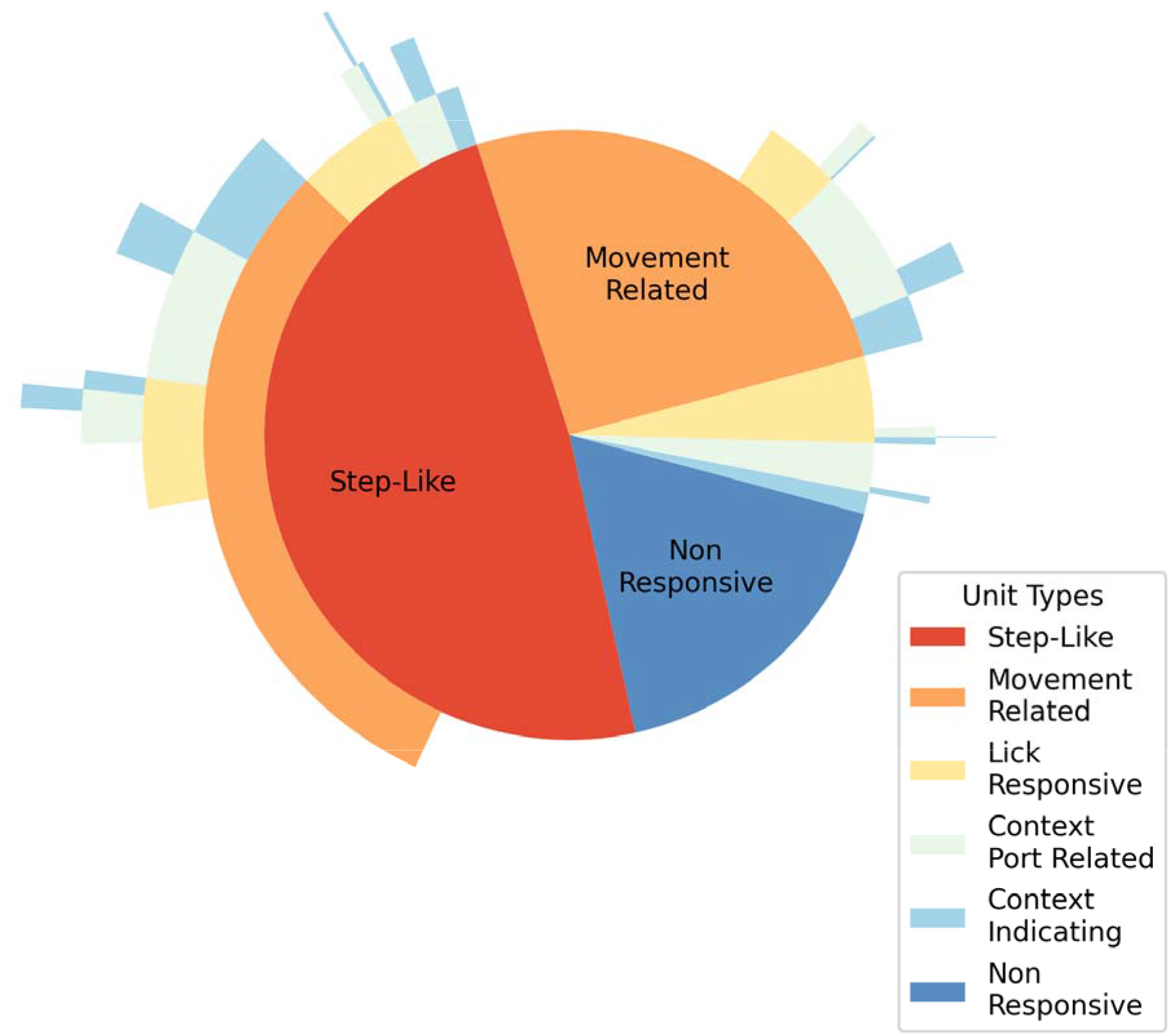
Classification and overlap of DMS unit response types during patch-foraging behavior. Sunburst diagram showing the distribution of response types among all recorded DMS units (n = 1,849). Step-like units exhibit detectable step transitions in firing rate during the post-reward period in the time-investment port. Movement-related units show modulated activity during locomotion into or out of either port, which could be for general travel or port-specific entry/exit responses. Lick-responsive units display firing modulation phase-locked to the lick cycle during continuous licking behavior throughout port occupancy. Context port-related units exhibit activity modulation during the fixed inter-reward intervals in the context port or responses to the depletion cue. Context- indicating units modulate their firing rate during the initial 1.25 seconds following context port entry, before reward delivery, based on the expected reward-rate block from the previous trial. Non-responsive units show no detectable modulation to any measured task variables.

**Figure 2–figure supplement 3:**
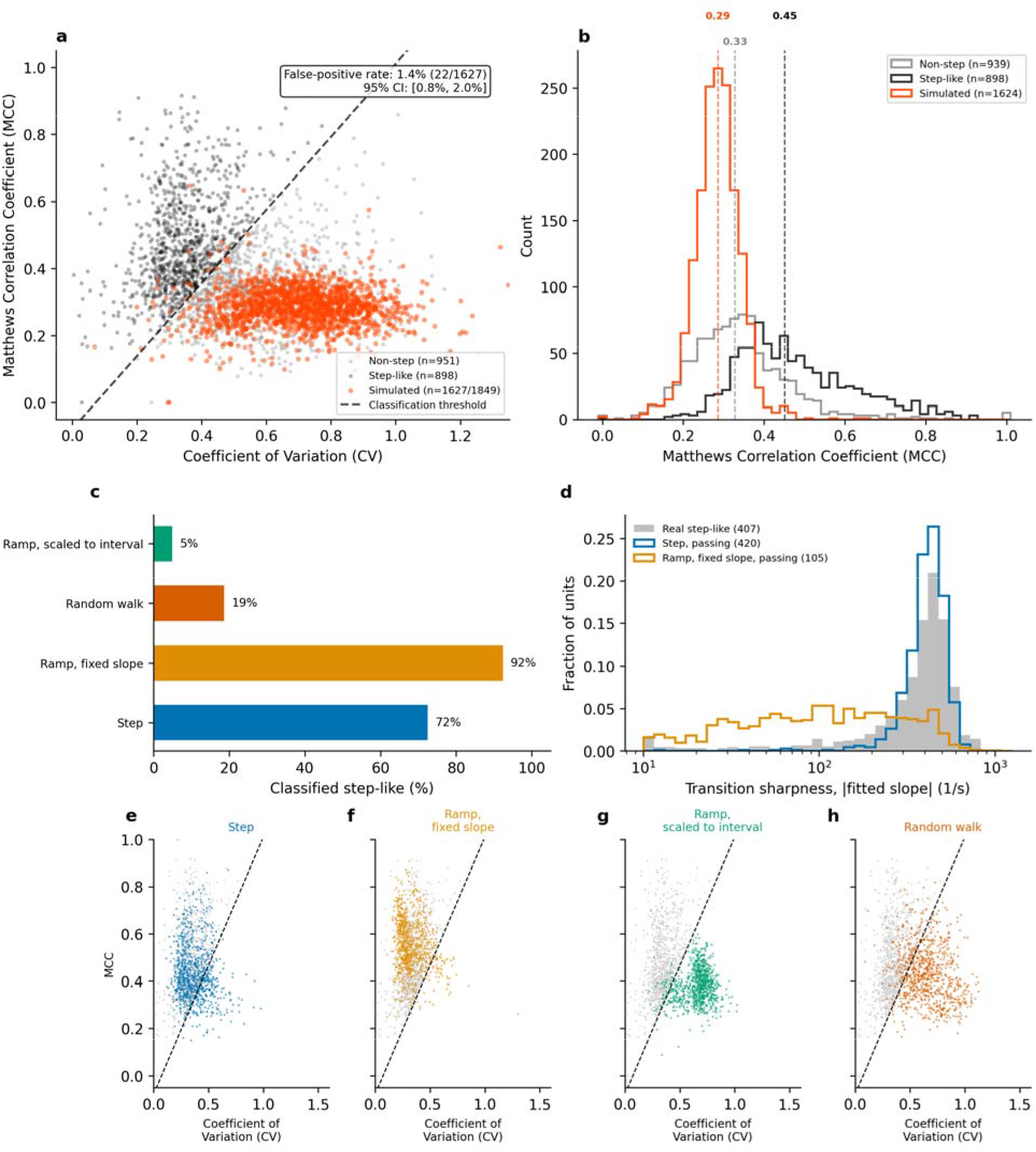
Step detection applied to simulated neurons with flat, ramping, random-walk and stepping firing rates. **a)** SVM classification scatter plot showing real step-like units (black, n = 898), real non- step units (gray, n = 951), and simulated flat-rate Poisson neurons (orange-red, n = 1,627) in CV-MCC feature space. The dashed line indicates the linear SVM decision boundary. Simulated neurons cluster below the boundary in the low-MCC, high-CV region. False positive rate: 1.35% (22/1,627; 95% CI [0.85%, 2.04%]). **b)** Step-style histogram comparing MCC distributions across the three populations. Median MCC values: simulated = 0.29, real non-step = 0.33, real step-like = 0.45 (Mann-Whitney U, simulated vs. step-like: p = 2.97 × 10 ² ³). For each of the 1,849 recorded units, a matched homogeneous Poisson process neuron was generated with the same mean firing rate and reward interval structure but zero modulation. Simulated spike trains were processed through the identical classification pipeline (sigmoid fitting, CV and MCC computation, SVM classification). Of 1,849 simulated units, 222 were excluded at the sigmoid fitting stage due to insufficient valid intervals (predominantly very low firing rate units, median 0.42 Hz). **c-h)** Simulated units with continuous rate changes, matched to each real step-like unit’s firing rates and interval structure at the modulation strength matched to the real units, passed through the same pipeline (Methods). **c)** Percentage classified as step-like for ramps scaled to the interval (5%), random walks (19%), ramps with a fixed slope (92%) and steps (72%). **d)** Transition sharpness (absolute fitted slope) of real step-like units and of the simulated steps and fixed-slope ramps that were classified step-like (medians 407, 420 and 105 per second). **e-h)** Each simulated kind on the classification plane of a, with real step-like units in gray and the classification boundary dashed.

**Figure 2–figure supplement 4.**
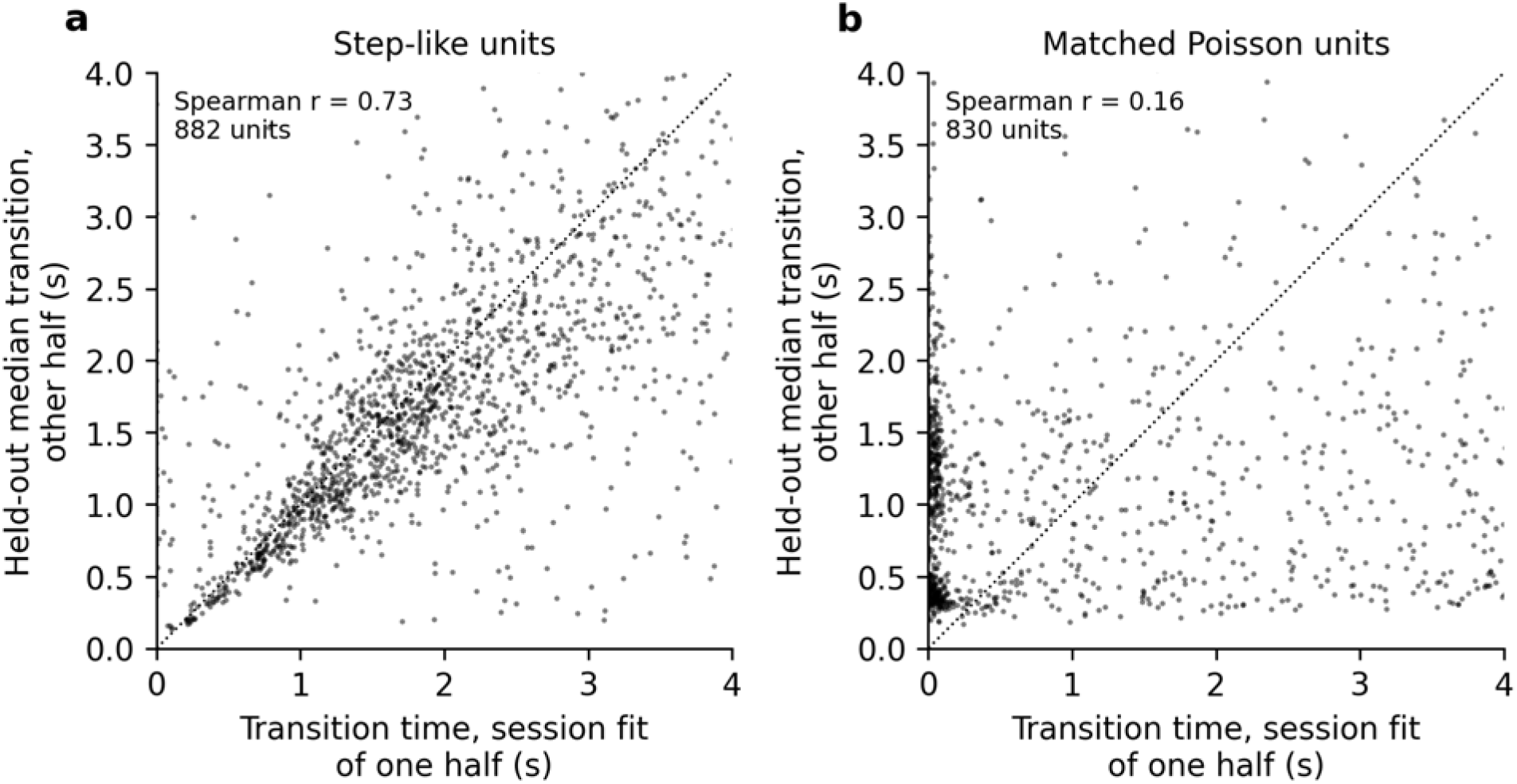
Transition times are consistent in held-out data. For each unit, the reward intervals were split in half at random. The transition time from the session-level fit of one half (x axis) is plotted against the median transition time of the other, held-out half (y axis), whose intervals were fit under the first half’s bounds, each starting from the held-out half’s own session-level fit. Both directions are shown. **a)** Step-like units (Spearman r = 0.73, 882 units). **b)** Simulated flat-rate Poisson units matched to each step-like unit’s firing rate and intervals (Spearman r = 0.16, 830 units). Points along the left edge are halves whose session fit found no transition and placed it at the start of the interval.

**Figure 3–figure supplement 1:**
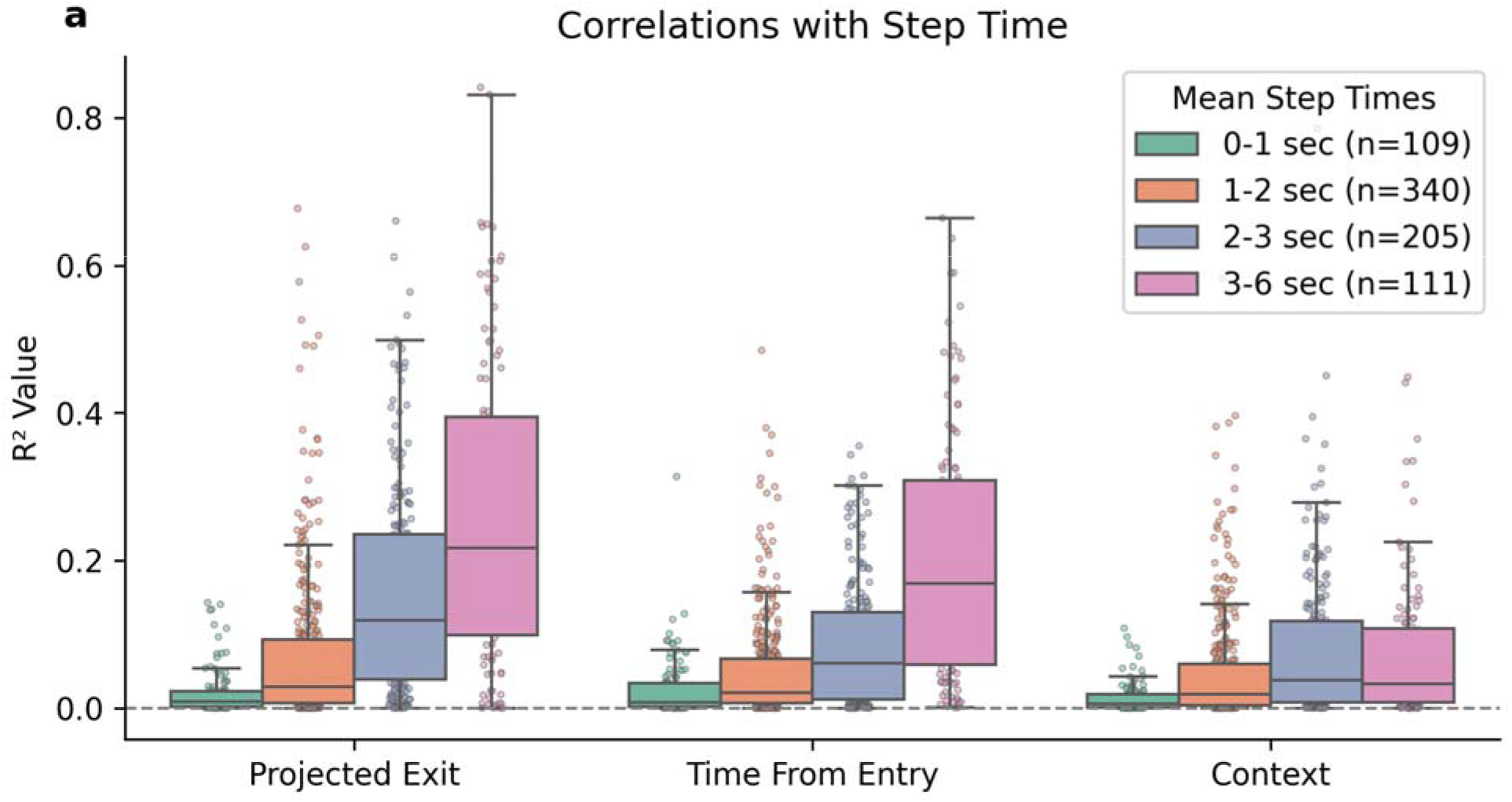
Correlation strength between neural step times and task variables varies by mean step time of DMS units. Box plots showing the distribution of R² values quantifying the relationship between individual unit step times and three task variables: projected exit time (time the mouse would wait following each reward based on session-specific behavioral models), time of reward relative to port entry, and context block reward rate. Units are grouped by their mean step time. Individual R² values for each unit are overlaid as points using a swarm plot distribution to avoid overlap.

**Figure 3–figure supplement 2.**
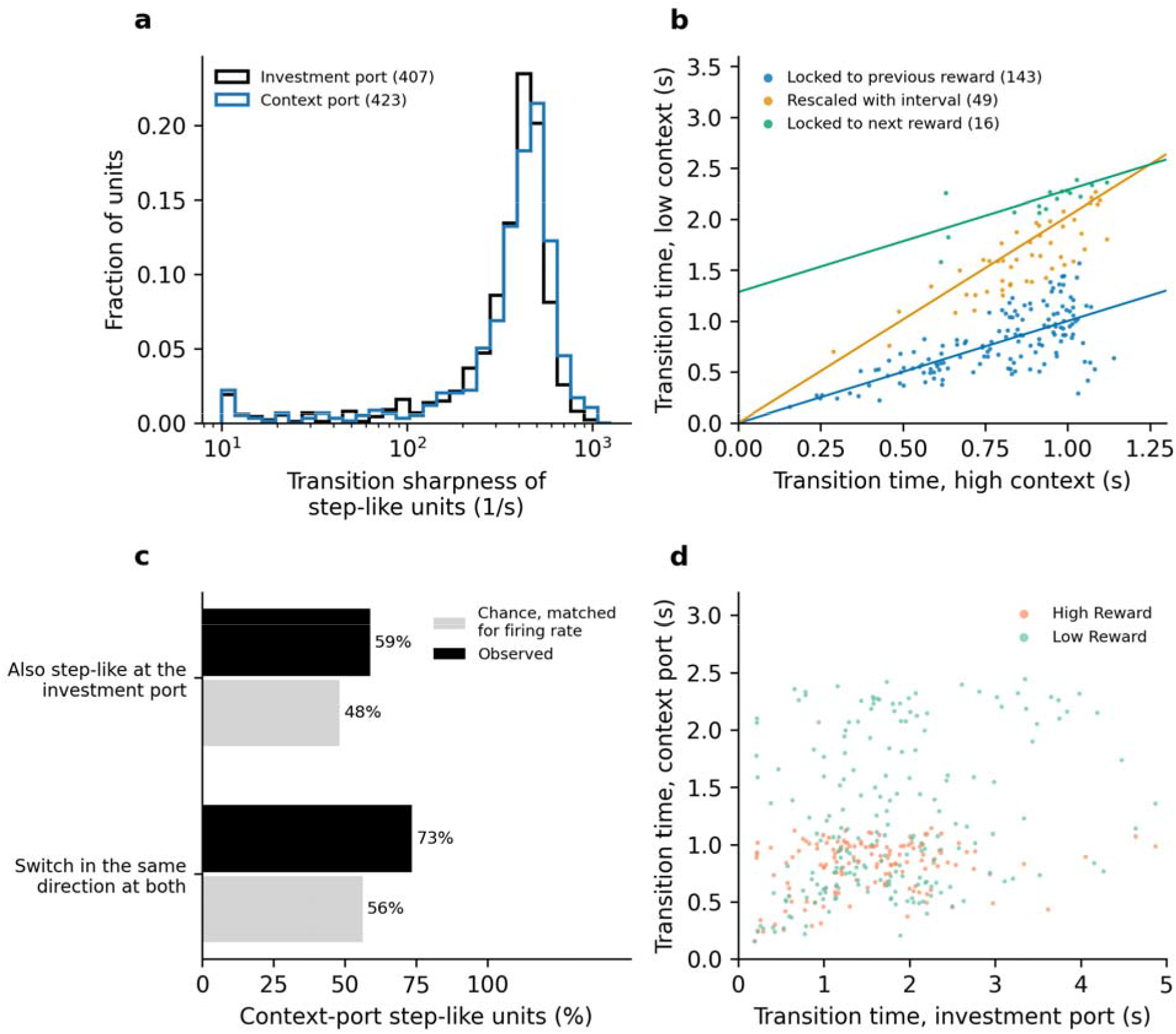
Step-like activity at the context port. The step detection and classification of Figure 2 applied to the context port’s reward intervals, with the same classification boundary. Of 1,890 context-port units, 601 were step-like (each port’s analysis includes the units with a usable interval at that port, so the total differs from Figure 2’s 1,849). **a)** Transition sharpness of step-like units at the time-investment and context ports (medians in parentheses). **b)** For each context-port step-like unit with at least five valid transitions in each context (208 units), the median transition time after a reward in the low reward-rate context against the high reward-rate context. Each unit is colored by the prediction it is closest to. The three predictions are transitions locked to the previous reward (identity line), transitions that rescale with the interval (ratio 2.03, the ratio of the two contexts’ inter-reward intervals), and transitions locked to the next reward. Across units, transitions were 1.29 times later in the low reward-rate context (95% CI 1.22 to 1.36, linear mixed model). **c)** Of the 577 context-port step-like units also analyzed at the time- investment port, the percentage also step-like there, and of those, the percentage that switch in the same direction at both ports, against chance matched for session firing rate. **d)** Transition times of the same units at the two ports, by context.

**Figure 4–figure supplement 1.**
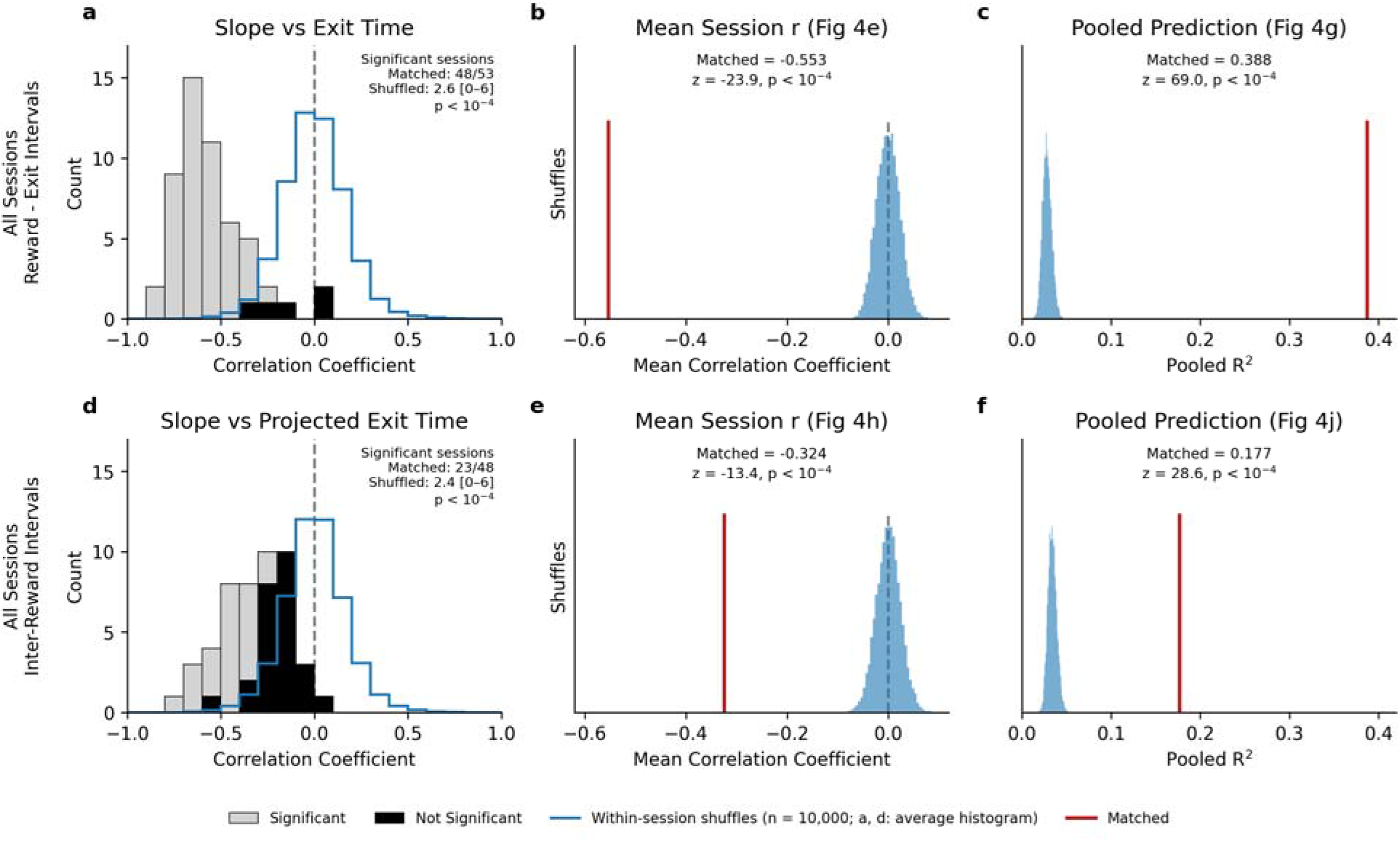
The accumulation-based exit prediction depends on the trial-by-trial pairing of neural activity and behavior. Within each session, accumulation traces (fitted slope and intercept together) were randomly reassigned among intervals of the same type, with behavior held fixed (10,000 shuffles). **a-c)** Reward-to- exit intervals (as in Figure 4e–g, 53 sessions, 2,410 intervals). **d-f)** Inter-reward intervals (as in Figure 4h–j, 48 sessions with at least 10 intervals, 2,622 intervals). **a, d)** Per-session Pearson correlation between accumulation slope and exit time (a) or projected exit time (d). Light gray, sessions with p < 0.05 (two-sided). Black, not significant. Blue outline, the per-session correlation histogram under shuffling, averaged over all shuffles. **b, e)** Distribution of the mean per-session correlation across shuffles (blue), against the matched value (red). **c, f)** Distribution of the pooled R² between neurally predicted and true (c) or projected (f) exit time across shuffles (blue), against the matched value (red). No shuffle reached the matched value for any statistic (permutation p < 10 ). z = (matched − shuffle mean) / shuffle standard deviation.

**Figure 4–figure supplement 2.**
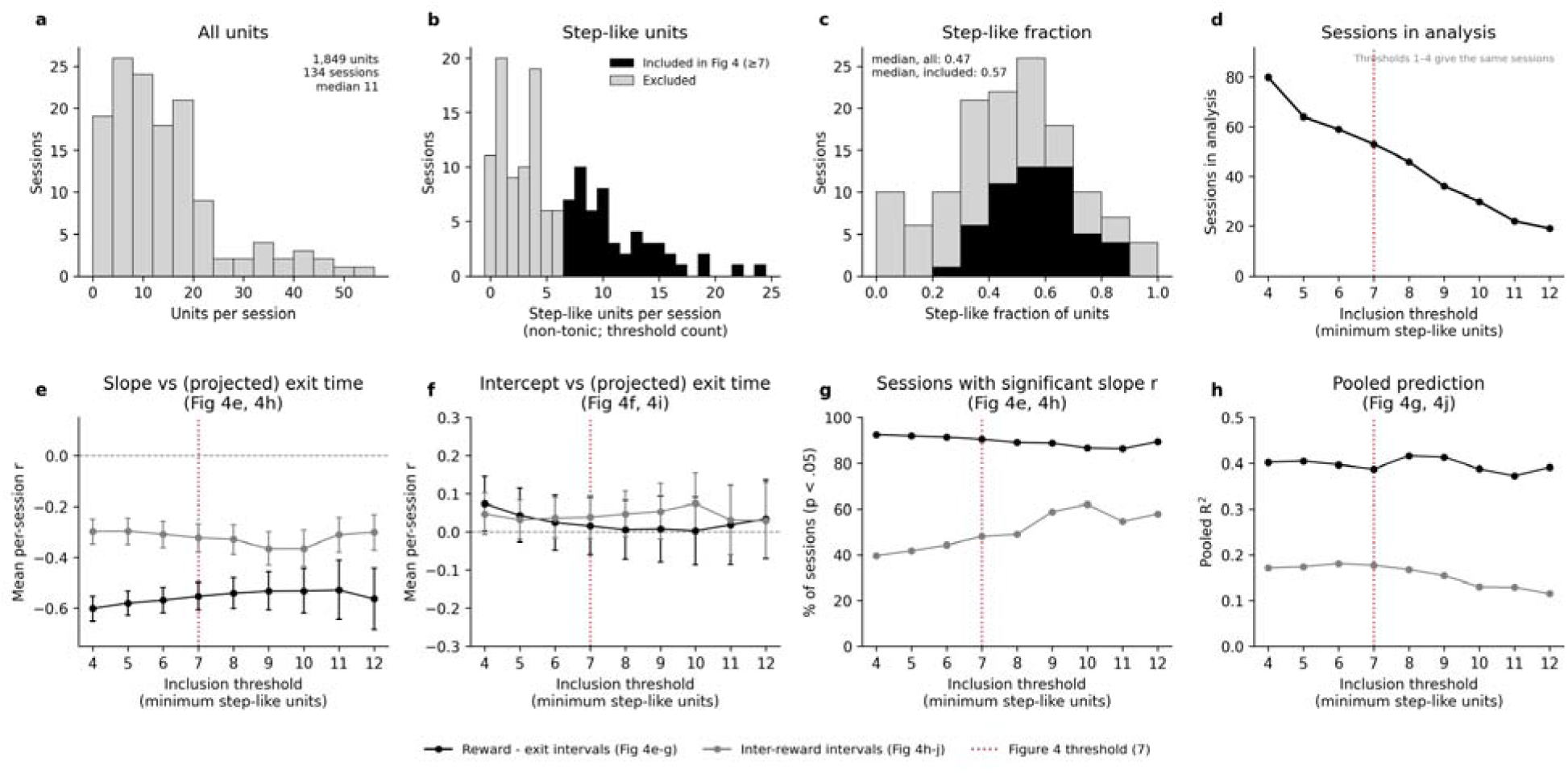
Accumulation results are qualitatively consistent across session inclusion thresholds. **a)** Units per session (1,849 units in 134 sessions, median 11 per session). Units were counted if they had more than 1,000 spikes and at least one usable interval with a fitted transition. **b)** Step-like units per session, as counted for session inclusion. Black, sessions meeting the Figure 4 threshold (at least 7). Gray, excluded. **c)** Fraction of units per session that were step-like, shaded as in b. **d)** Sessions entering the analysis at each inclusion threshold. Thresholds of 4 or fewer are equivalent, because fitting an interval’s accumulation line requires at least five step positions. **e-h)** Figure 4 statistics recomputed at each threshold, for reward-to-exit intervals (black, as in Figure 4e–g) and inter-reward intervals (gray, as in Figure 4h–j). **e)** Mean per-session correlation between accumulation slope and exit time (or projected exit time), ±95% confidence interval across sessions. **f)** As e, for the intercept. **g)** Percentage of sessions with a significant slope correlation (p < 0.05, two-sided). **h)** Pooled R² between neurally predicted and true (or projected) exit time. Dotted line, the threshold used in Figure 4 (7).

**Figure 4–figure supplement 3.**
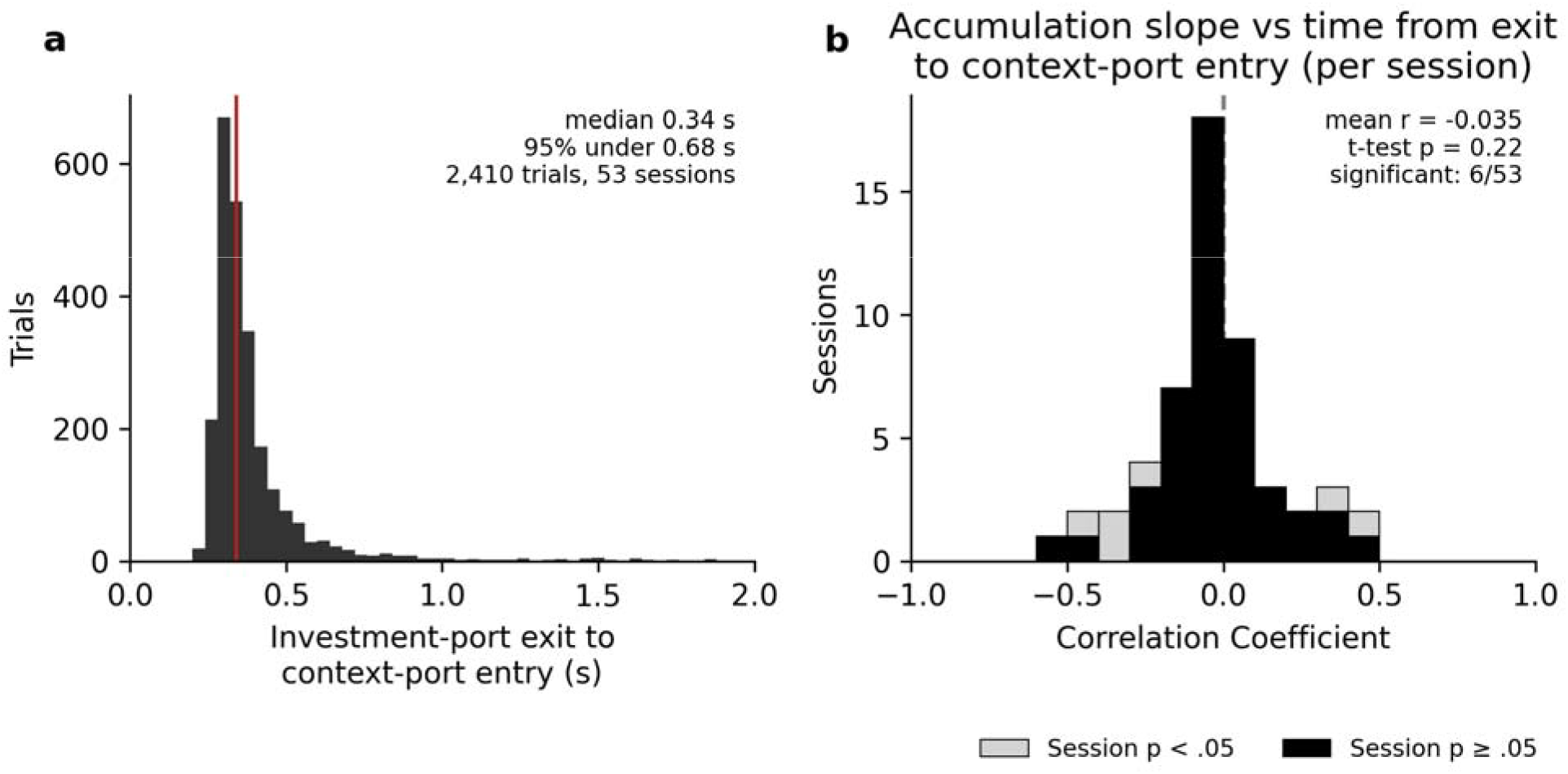
Time from investment-port exit to context-port entry, and the accumulation slope. **a)** Distribution of the time from exit (the final withdrawal from the time-investment port before the context-port visit) to the next context-port entry, for the reward-to-exit intervals of Figure 4e (2,410 trials, 53 sessions, median 0.34 s, red line). **b)** Per session, the Pearson correlation between the accumulation slope on the final-reward-to-exit interval and that time. Sessions as in Figure 4e. Light gray, p < 0.05. Mean r = −0.035 (one-sample t-test, p = 0.22), 6 of 53 sessions individually significant.

